# A regulatory role of novel long non-coding RNA, *BAZ1A-AS1*, in vascular smooth muscle cell functions during neointima proliferation in human saphenous veins

**DOI:** 10.64898/2026.02.03.703576

**Authors:** David S. Kim, Brandee Goo, Praneet Veerapaneni, Ronnie Chouhaita, Medha Guduru, Nicole Cyriac, Mourad Ogbi, David J. Fulton, Wei Zhang, Xiaochun Long, Avirup Guha, Philip Coffey, Robert D. Rice, Dominic R. Gallo, Vijay S. Patel, Richard Lee, Ha Won Kim, Hong Shi, Neal L. Weintraub

## Abstract

**Background:** Neointimal proliferation (NP) is a major cause of stenosis and occlusion in arteries and veins. Human saphenous veins (HSV), often used for coronary artery bypass grafting, frequently fail in part due to NP. Long non-coding RNAs (lncRNAs) have emerged as critical regulators of vascular smooth muscle cell (VSMC) phenotype, yet lncRNAs governing NP in human veins remain largely uncharacterized.

**Methods and Results:** Using an *ex vivo* model of NP in human SV, we performed bulk RNA sequencing on SV tissues with and without NP. Among differentially expressed lncRNAs, we identified a previously uncharacterized transcript, *Bromodomain Adjacent to Zinc Finger A1 antisense 1* (*BAZ1A-AS1*), and its predicted *cis*-regulatory partner gene *BAZ1A*, as markedly upregulated during NP and predominantly enriched in VSMCs. Both transcripts were induced by genotoxic stimuli, with *BAZ1A-AS1* showing transient induction preceding sustained *BAZ1A* upregulation, consistent with a priming role in the DNA damage response. Silencing of either *BAZ1A-AS1* or *BAZ1A* attenuated VSMC proliferation and migration, accompanied by upregulation of contractile markers and suppression of proliferative and inflammatory transcriptional programs. ChIRP-qPCR demonstrated that *BAZ1A-AS1* physically interacts with 3′ distal exons of *BAZ1A* at the DNA level while selectively engaging *CNN1*, *CCND1*, and *IL6* mRNAs, indicating mechanistically distinct chromatin- and RNA-level regulatory functions. *Baz1a* haploinsufficiency attenuated neointima formation and preserved VSMC contractile identity in a mouse carotid artery ligation model.

**Conclusions:** We identify *BAZ1A-AS1* and *BAZ1A* as stress-responsive regulators of VSMC phenotype, operating through dual mechanisms of cis-regulatory chromatin interaction and selective mRNA engagement, and demonstrate a novel role for the *BAZ1A*-*AS1*/*BAZ1A* axis in promoting NP.

## Introduction

Neointimal proliferation (NP) is a major cause of stenosis in both arterial and venous vascular disease. In the venous circulation, NP is particularly relevant to the human saphenous vein (HSV), which remains the most widely used bypass conduit for coronary artery bypass grafting and is susceptible to progressive graft narrowing and failure.^1–3^ Despite the clinical burden, no pharmacological strategy has proven effective at preventing NP, reflecting an incomplete understanding of its molecular basis.

NP is driven by injury-induced phenotypic switching of vascular smooth muscle cells (VSMCs) from a quiescent, contractile to a proliferative, synthetic state.^4–6^ While signaling pathways that promote VSMC dedifferentiation have been extensively studied in rodent arterial injury models, the regulatory mechanisms governing this process in human venous tissue remain poorly defined. This gap is significant because species-specific regulatory elements, particularly those incorporated in the noncoding genome, are unlikely to be captured by rodent-based discovery.

Long noncoding RNAs (lncRNAs) represent human-specific epigenetic regulators that exemplify this challenge. LncRNAs, which frequently act *in cis* to modulate neighboring protein-coding genes, have established roles in VSMC phenotypic control: *CARMN* maintains contractile identity by activating myocardin,^7,8^ whereas *INKILN* promotes pro-inflammatory gene expression through scaffolding of *MKL1*.^9^ LncRNAs dysregulated during NP in human vascular tissue have not been systematically characterized. Furthermore, because lncRNAs are poorly conserved across species, rodent-based discovery is unlikely to fully capture the lncRNA networks governing human vascular disease.

To address this gap, we employed bulk RNA sequencing and spatial transcriptomics in an *ex vivo* model of NP in HSV to identify lncRNAs associated with this process. We identified a previously uncharacterized lncRNA, *Bromodomain Adjacent to Zinc Finger A1 antisense 1* (*BAZ1A-AS1*), and its predicted *cis*-regulatory partner *BAZ1A*, a chromatin remodeling factor with established roles in cellular stress responses, as coordinately induced in VSMCs during NP. Functional studies demonstrated that knockdown of this lncRNA-gene pair suppressed VSMC proliferation, migration, and synthetic gene expression, while *in vivo* haploinsufficiency of *Baz1a* in mice attenuated neointima formation and preserved contractile identity following vascular injury. These findings identify the BAZ1A-AS1/*BAZ1A* axis as a previously unrecognized regulator of VSMC phenotype switching and a potential contributor to the pathogenesis of obstructive vascular disease.

## Methods

### Human SV ex vivo culture

The method for collecting and culturing human saphenous vein (HSV) was adapted from our previous protocols.^10,11^ Briefly, freshly obtained SV samples from patients undergoing coronary artery bypass grafting (CABG) were transported directly to the laboratory. The HSV segments were rinsed in sterile phosphate-buffered saline (PBS) and transversely sectioned into 3 mm pieces; some were stored at -80°C for later analysis (Day 0). The remaining segments were cultured for 7 days (Day 7) in RPMI 1640 medium supplemented with 2 g/L NaHCO3, 100 IU/ml penicillin, 100 µg/ml streptomycin, 4 ml/L L-glutamine, and 30% fetal bovine serum (FBS), incubated at 37°C in a 5% CO2 atmosphere. The culture medium was refreshed every other day, after which the segments were stored at -80°C for future analyses. The collection of HSV samples and associated experiments were exempted from human subjects research requirements by the Institutional Review Board (IRB) at Augusta University. All samples were collected using the endovascular harvesting (EVH) technique. Clinical characteristics of patients whose HSV were used for bulk RNA sequencing are provided in **Supplemental Table S1.**

### Bulk RNA sequencing

Bulk RNA sequencing of Day 0 and Day 7 whole HSV tissue was performed as previously described.^11^ For knockdown transcriptomic studies, total RNA from HSVSMCs (for *BAZ1A-AS1* knockdown) and RD cells (for *BAZ1A* knockdown under UV exposure) with RNA integrity number (RIN) greater than 8 and passed downstream library QC was sent to Genewiz or Genomics Core at Augusta University for preparation into RNA-Seq libraries. Polyadenylated RNA or ribosome-depleted RNA was sequenced at a depth of 20 million reads per replicate using Illumina HiSeq 2500 system, using a 150 bp paired-end protocol (Illumina). Raw reads were trimmed using Cutadapt version 4.1, and the processed reads were mapped to the GRCh38/hg38 genome assembly by using STAR version 2.7.0.^12^ DESeq2 version 1.22 was used to generate normalized read counts and conduct differential gene expression analysis with a false discovery rate (FDR) threshold of less than 0.05 using R studio version 494 (https://posit.co/download/rstudio-desktop/) and public server of usegalaxy.org.^13,14^ Principal component analysis (PCA) and volcano plot was created through ggplot2. Gene ontology analysis and gene-network graph were performed and created through clusterProfiler version 3.0.4.^15^

### Hallmark gene set enrichment analysis

Hallmark gene set enrichment analysis (GSEA) was performed using the fgsea package in R with gene sets from the Molecular Signatures Database (MSigDB) Hallmark collection. Genes were ranked by the Wald statistic from DESeq2 differential expression analysis. Enrichment was considered significant at FDR < 0.05 after Benjamini-Hochberg correction. For convergence analysis, normalized enrichment scores (NES) from *BAZ1A-AS1* knockdown (HSVSMCs) and *BAZ1A* knockdown (RD cells under UV) were plotted against each other for all Hallmark pathways.

### Hi-C data visualization

Published Hi-C contact data from human aorta were obtained from GEO (GSE58752).^16^ Contact frequency maps within a 500-kilobase window centered on the *BAZ1A-AS1/BAZ1A* locus on chromosome 14 were visualized to identify chromatin interaction partners.

### Visium spatial transcriptomics

The method for analysis of visium spatial transcriptomics was adapted from our previous protocols^10,11,17^. Formalin-fixed paraffin-embedded (FFPE) Visium Spatial Transcriptomics data were generated following the manufacturer’s protocol. Briefly, 5-µm FFPE sections from three biological replicates of Day 0 and Day 7 HSV samples with a DV200 above 50% were mounted on glass slides and processed at the University of Michigan’s Advanced Genomics Core. After H&E imaging, probe hybridization, ligation, extension, and library construction were carried out using the Visium FFPE Gene Expression workflow. cDNA libraries were sequenced on an Illumina NovaSeq 6000. Demultiplexed FASTQ files were converted to count matrices using SpaceRanger v1.3.1 (10X Genomics) and mapped to GRCh38. Processed data utilized the Visium Human Transcriptome Probe Set v2.0 with spot annotations performed in Loupe Browser 5. Seurat Objects were constructed from count matrices and spot annotations, excluding spots with fewer than 500 UMIs and over 5% mitochondrial reads as low-quality. SCTransform normalized high-quality spots, which were pooled, reduced in dimensionality, and clustered (∼5000 spots) using Seurat. Cellular identities for each cluster were predicted using published single-cell transcriptomic signatures and tissue morphology of human HSV samples. Differentially expressed genes (DEGs) with an FDR < 0.05 were identified using the Benjamini-Hochberg method, with top DEGs selected based on absolute values.

### *In silico* coding potential analysis

The coding potential of *BAZ1A-AS1* was assessed using five complementary approaches: PhyloCSF (codon substitution frequency across species), CPAT (coding probability based on ORF size, coverage, and hexamer usage), Lee translational initiation site prediction, PRIDE reprocessing (mass spectrometry database search for translated peptides), and Bazzini small ORF detection. All metrics uniformly classified *BAZ1A-AS1* as non-coding.

### 3’ Rapid Amplification of cDNA Ends (RACE)

3’ RACE was performed on total RNA isolated from HSVSMCs using the 3’ RACE Systems for Rapid Amplification of cDNA Ends (Invitrogen) according to the manufacturer’s protocol. Gene-specific primers complementary to *BAZ1A-AS1* were used to amplify the 3’ end. Products were resolved by agarose gel electrophoresis, gel-purified, cloned, and Sanger-sequenced to confirm a single 2,116-nucleotide isoform.

### Quantitative PCR

Total RNA was extracted from HSV or cultured cells with QIAzol Lysis Reagent, and purified with miRNeasy Mini Kit (Qiagen). Real time quantification of mRNA levels of the genes of interest was performed using Brilliant II SYBR Green QPCR Master Mix (Agilent Technologies) per manufacturer’s instructions. Normalized Ct values were subjected to statistical analysis and fold change was calculated by ΔΔ Ct method as described previously, normalized against 18S ^18,19^. Primer sequences are listed in **Supplemental Table S2**.

### Isolation of human saphenous vein smooth muscle cells (HSVSMCs)

Saphenous vein segments were rinsed with sterile PBS, the adventitia was carefully removed, and the vein was cut longitudinally and pinned. The intimal layer was gently scraped with a surgical scalpel to remove endothelial cells. The medial layer was peeled and minced into 1 mm^2^ pieces before being plated on a 12-well plate at 37°C in Smooth Muscle Cell Growth Medium (SmGm-2, Lonza) with 20% fetal bovine serum and 1% penicillin-streptomycin.

### Isolation of mouse aortic vascular smooth muscle cells (MASMCs)

Aortas were harvested from *Ldlr^-/-^* and *Baz1a^+/-^;Ldlr^-/-^* mice, cleaned of adventitial tissue, and digested enzymatically in collagenase/elastase solution. Isolated cells were cultured in Dulbecco’s Modified Eagle Medium (DMEM) supplemented with 10% FBS and 1% penicillin-streptomycin at 37°C in a 5% CO₂ humidified incubator. Experiments were performed between passages 2 and 5.

### Culture and transfection of HSVSMCs, HCASMCs, and rhabdomyoblasts (RDs)

HSVSMCs and human coronary artery smooth muscle cells (HCASMCs) were cultured in DMEM or SmGm-2 supplemented with 10% FBS and 1% penicillin-streptomycin at 37°C in a 5% CO₂ humidified incubator. RDs were cultured in DMEM supplemented with 10% FBS and 1% penicillin-streptomycin at 37°C in a 5% CO₂ humidified incubator. RD cells were used for *BAZ1A* knockdown experiments requiring sustained culture, as *BAZ1A s*ilencing caused significant cytotoxicity in primary HSVSMCs that precluded long-term downstream analysis. Cells were seeded in 6-well plates and grown to 60-70% confluence before starvation overnight with 0.2% FBS. For *BAZ1A-AS1* knockdown, cells were transfected with 25 nM locked nucleic acid (LNA) gapmeR antisense oligonucleotides (ASO; Qiagen), which preferentially degrade nuclear transcripts. For *BAZ1A* knockdown, cells were transfected with 25 nM siRNA (Qiagen) targeting mature cytoplasmic mRNA. All transfections were performed using Lipofectamine RNAiMAX (Invitrogen) according to the manufacturer’s protocol. Cells were collected 24-48 hours post-transfection.

### UV exposure

HSVSMCs and RD cells were exposed to ultraviolet C (UVC, 254 nm) irradiation using the germicidal lamp of a biological safety cabinet (BSC). Culture medium was removed and cells were washed once with PBS immediately before exposure. With the sash positioned at working height, cells were irradiated for 5, 15, or 30 seconds. Fresh culture medium was added immediately following irradiation and cells were returned to the incubator. For time-course experiments, cells were harvested at 0, 2, 6, 12, 18, and 24 hours post-exposure.

### Conditioned media preparation

Conditioned media (CM) was collected from Day 7 HSV explant cultures. After 7 days of culture, supernatant was collected, centrifuged at 300 x g for 5 minutes to remove debris, and filtered through a 0.45 µm filter. CM was applied directly to HSVSMCs for 24 hours before RNA extraction.

### Fluorescent In Situ Hybridization

Samples were fixed with 4% paraformaldehyde and permeabilized with 0.1% Triton X-100. Fluorescently labeled FISH probes (Biosearch Technologies) specific to *MALAT1* and *BAZ1A-AS1* were hybridized at 37°C for 12-16 hours. Post-hybridization, samples were washed in saline-sodium citrate (SSC) buffer to remove excess probes, then mounted with DAPI for nuclear staining. Images were captured using a fluorescence microscope, and RNA localization was analyzed using ImageJ software.

### Nuclear and Cytoplasmic RNA separation

Cells were harvested and washed with cold phosphate-buffered saline (PBS), followed by lysis in a hypotonic buffer to disrupt the plasma membrane. The lysate was centrifuged at 1,000 x g for 5 minutes at 4°C to pellet the nuclei, separating the cytoplasmic fraction in the supernatant. The nuclear pellet was washed and further lysed in nuclear isolation buffer. RNA from both the nuclear and cytoplasmic fractions was extracted using Rneasy Mini Kit (Qiagen), following the manufacturer’s protocol. The quality and purity of the isolated RNA were assessed by spectrophotometry and agarose gel electrophoresis.

### EdU-Amplex Red proliferation assay

Cell proliferation was assessed using the Click-iT EdU Microplate Assay with Amplex Red detection (Invitrogen) according to the manufacturer’s instructions. Cells were incubated with 10 µM EdU for 18–24 hours. After fixation and permeabilization, incorporated EdU was detected via copper-catalyzed click chemistry and Amplex Red fluorogenic substrate. Fluorescence was measured using a microplate reader at excitation/emission of 530/590 nm. Results were normalized to non-transfected controls.

### CCK-8 viability assay

Cell viability was assessed using the Cell Counting Kit-8 (CCK-8; Dojindo) according to the manufacturer’s protocol. Cells were incubated with 10 µL CCK-8 reagent per 100 µL culture medium for 1–2 hours at 37°C. Absorbance was measured at 450 nm using a microplate reader. Results were normalized to vehicle-treated controls.

### LDH release assay

LDH release was measured using the CyQUANT LDH Cytotoxicity Assay (Millipore Sigma) according to the manufacturer’s instructions. Culture supernatants were collected 24 hours post-transfection. Spontaneous LDH release from medium-only controls was subtracted from experimental values. Absorbance was measured at 490 nm with background subtraction at 680 nm.

### Wound scratch assay

HSVSMCs were seeded in 6-well plates and grown to confluence. After overnight serum starvation (0.2% FBS), a uniform scratch was created using a sterile 200 µL pipette tip. Detached cells were removed by washing with PBS, and cells were maintained in 0.2% FBS medium. Brightfield images were captured at 0, 12, and 24 hours using an inverted microscope. Wound closure was quantified as the percentage of area closed relative to the initial wound area using ImageJ software.

### Annexin V and Propidium Iodide Staining

Cells were collected and washed with cold phosphate-buffered saline (PBS), then resuspended in binding buffer at a concentration of 1 × 10^6^ cells/mL. Annexin V conjugated to a fluorescent dye and propidium iodide (PI) were added according to the manufacturer’s instructions (ThermoFisher, V13242). The cell suspension was incubated in the dark for 15 minutes at room temperature to allow staining. For flow cytometry-based analysis, samples were analyzed within 1 hour to detect apoptosis. Annexin V positivity indicated early apoptosis, while dual Annexin V and PI staining indicated late apoptosis or necrosis. For immunofluorescence-based analysis under basal and UV-exposed conditions, stained cells were imaged directly on coverslips and PI fluorescence intensity was quantified using ImageJ software.

### H3 and H4 histone modification profiling of human SV

Histone modification profiling was conducted using the EpiQuik™ H3/H4 Modification Multiplex Assay Kit (EpigenTek, P-3102-96). Nuclear extracts were prepared from Day 0 and Day 7 HSV tissue according to the kit’s protocol. The histone proteins were bound to the assay wells, followed by incubation with modification-specific antibodies to detect the specified H3 and H4 modifications. After washing, a secondary antibody was applied for colorimetric detection, and the absorbance was measured at 450 nm using a microplate reader. Data were analyzed to quantify the levels of each histone modification.

### Chromatin Isolation by RNA Purification (ChIRP)-qPCR

ChIRP-qPCR was performed as described previously.^20^ In short, 2 × 10^6^ HSVSMCs were crosslinked with 1% glutaraldehyde for 15 minutes, and quenched with 1.25 M glycine. Cells were pelleted at 2000 RCF for 5 min at 4°C. Biotinylated antisense oligonucleotide probes (Biosearch Technologies) complementary to *BAZ1A-AS1,* excluding the region overlapping with *BAZ1A* exon 1, were hybridized to the crosslinked chromatin. Pooled probes were divided into either even and odd pools based on their sequential order, with every other probe assigned to a different pool to serve as internal controls. The complexes were then captured using streptavidin-coated magnetic beads (Invitrogen). After washing, the bound material was eluted, and DNA was purified using DNA Clean & Concentrator Kit (Zymo Research) and RNA was purified using the Direct-zol RNA MiniPrep Kit (Zymo Research). qPCR was performed on the eluted DNA to assess physical binding to genomic loci (*BAZ1A* exons, *NFKBIA*, *SRP54*) and on eluted RNA to assess mRNA interactions (*CNN1*, *CCND1*, *IL6*, *BAZ1A*, *CDK4*, *CDK6*, *ACTA2*, *NFKBIA*). Enrichment for each target was calculated relative to *GAPDH* as a negative control. Even and Odd probe pools were compared to their respective same-pool *GAPDH* controls separately (not pooled). A target was considered enriched only if significant in both Even and Odd probe pools. Gel electrophoresis of ChIRP-qPCR products was performed after 28 cycles to visualize specific amplification. A target was considered enriched only if significant in both Even and Odd probe pools. Probe sequences and pool assignments are listed in **Supplemental Table S3**.

### Carotid artery ligation

Complete ligation of the left common carotid artery was performed in 8-12-week-old mice in either wild type (WT, C57BL/6) or low density lipoprotein receptor knockout (Ldlr-/-) background to induce neointima formation. Global *Baz1a* knockout mice were obtained from Dr. Scott Keeney (Memorial Sloan Kettering Cancer Center).^21^ Heterozygous knockout mice were used in this study, due to infertility exhibited by *Baz1a* homozygous knockout mice. Both male and female mice were used. Mice were anesthetized with isoflurane. The left carotid artery was ligated near the bifurcation with 6-0 silk, while the right carotid artery served as an uninjured contralateral control. After 21 or 28 days, mice were euthanized, and both carotid arteries were harvested and processed for histological analysis. H&E staining was used to assess neointima formation at 200 μm and 400 μm proximal to the ligation site, and intimal and medial areas were quantified using ImageJ software.

### Western blotting

Protein extraction and Western blotting were performed as described previously.^18^ Antibodies used are listed in **Supplemental Table S4**.

### Immunohistochemistry and histology

HSV tissues were fixed in 10% neutral buffered formalin for less than 24 hours, dehydrated with 70% ethanol followed by a series of graded alcohol immersions, cleared with xylene, and embedded in paraffin wax. Tissue processing, embedding, sectioning, and histological staining were performed by the Electron Microscopy and Histology Core Facility at Augusta University (RRID: SCR_026810). Paraffin-embedded HSV tissue sections (5 µm) were stained with hematoxylin and eosin (H&E), Verhoeff van Gieson (VVG), Masson’s trichrome, or processed for immunohistochemistry using antibodies against BAZ1A, αSMA, Ki67, or other targets as specified. DAB Substrate (Vector Labs) kits were used for visualization as previously described.^18^

For morphometric analysis, intima, media, adventitia, and total vessel area were quantified using ImageJ software with the freehand selection tool on VVG-stained slides, as previously described.^22^ The number of elastin breaks was counted in three sections per tissue sample to quantify elastin degradation. Elastin area fraction was quantified by thresholding VVG-stained sections in ImageJ, with the elastin-positive area expressed as a percentage of total vessel wall area within a defined region of interest.

For collagen quantification, Masson’s trichrome-stained sections were analyzed using the Color Deconvolution plugin in ImageJ (Masson Trichrome preset) to separate blue (collagen), red (muscle/cytoplasm), and background channels. On the blue channel, a consistent threshold was applied across all images to identify collagen-positive pixels. Collagen area fraction was calculated as the percentage of blue-stained area relative to total tissue area within the region of interest. Threshold settings were kept identical across all images, and all sections were photographed in one batch with matched white balance and exposure settings. For immunofluorescence quantification, Ki67-positive VSMCs were counted within the medial layer (delineated by internal and external elastic laminae) and expressed as a percentage of total DAPI-positive cells. αSma fluorescence intensity was quantified as mean fluorescence intensity within the vessel wall area using ImageJ. All quantification was performed by an observer blinded to genotype. Antibodies used are listed in **Supplemental Table S4.**

### Data Availability

Bulk RNA-seq data from Day 0 and Day 7 human saphenous veins (GSE310013) and Visium spatial transcriptomics data (GSE310351) were deposited in the Gene Expression Omnibus as part of our previous study.^11^ Bulk RNA-seq data from *BAZ1A-AS1* knockdown in HSVSMCs and *BAZ1A* knockdown in RD cells have been deposited in GEO under accession number GSE329766. All other data supporting the findings of this study are available from the corresponding author upon reasonable request.

### Statistical Analysis

All experiments were repeated at least 3 times independently, and statistical analysis was conducted with GraphPad Prism 10.0. Quantitative results are presented as mean ± SEM. In vivo data with sample size > 6 were tested for normal distribution by the Shapiro-Wilk test. A two-tailed unpaired t-test was performed when both groups were normally distributed, and the Mann-Whitney test was performed when one or more groups were not normally distributed. A t-test with Welch’s correction was performed for unpaired comparisons with unequal variances. A one-way ANOVA was performed for comparisons among more than two groups with equal variance, and the Brown-Forsythe test was used with unequal variance. A one-way ANOVA followed by Dunnett’s test was performed when two or more groups were compared with the same control (e.g., UV dose-response, Figure 2G). Two-way ANOVA followed by Tukey’s correction was used for multiple comparisons with two or more variables. For RNA-seq differential expression, Wald test with Benjamini-Hochberg correction was applied via DESeq2. For ChIRP-qPCR, Welch’s unpaired one-sided t-test (target > *GAPDH*) with Benjamini-Hochberg FDR correction across all target-pool comparisons was used (n = 3 biological replicates per pool). P < 0.05 was considered statistically significant. Details of statistical analysis are included in the figure legends for each individual experiment.

## Results

### *BAZ1A-AS1* as a novel lncRNA upregulated during neointima proliferation in human saphenous vein

Human saphenous vein (HSV) explants were obtained from patients undergoing CABG surgery and cultured for 7 days to induce NP (representative tissue morphology shown in **Figure 1K, L**).^10,11^ Bulk RNA-seq was performed on Day 0 and Day 7 HSV to identify differentially expressed (DE) lncRNAs. Clinical characteristics of patients whose HSV were used for bulk RNA sequencing are provided in **Supplemental Table S1**. Principal component analysis demonstrated clear separation between Day 0 and Day 7 transcriptomes (**Figure 1A**). We identified 2,429 DE lncRNAs between Day 0 and Day 7 HSV (|fold change| > 2.0, *P*-adj < 0.05), of which 1,186 were upregulated and 1,243 were downregulated at Day 7 (**Figure 1B**). Notably, lncRNAs previously reported to regulate cardiovascular function, including *NEAT1*, *CARMN*, *MALAT1*, *SNHG12*, *ACTA2*-*AS1*, and *H19*, were also DE, indicating that HSV engage established lncRNA regulatory networks during NP (**Figure 1C**).

**Figure 1.**
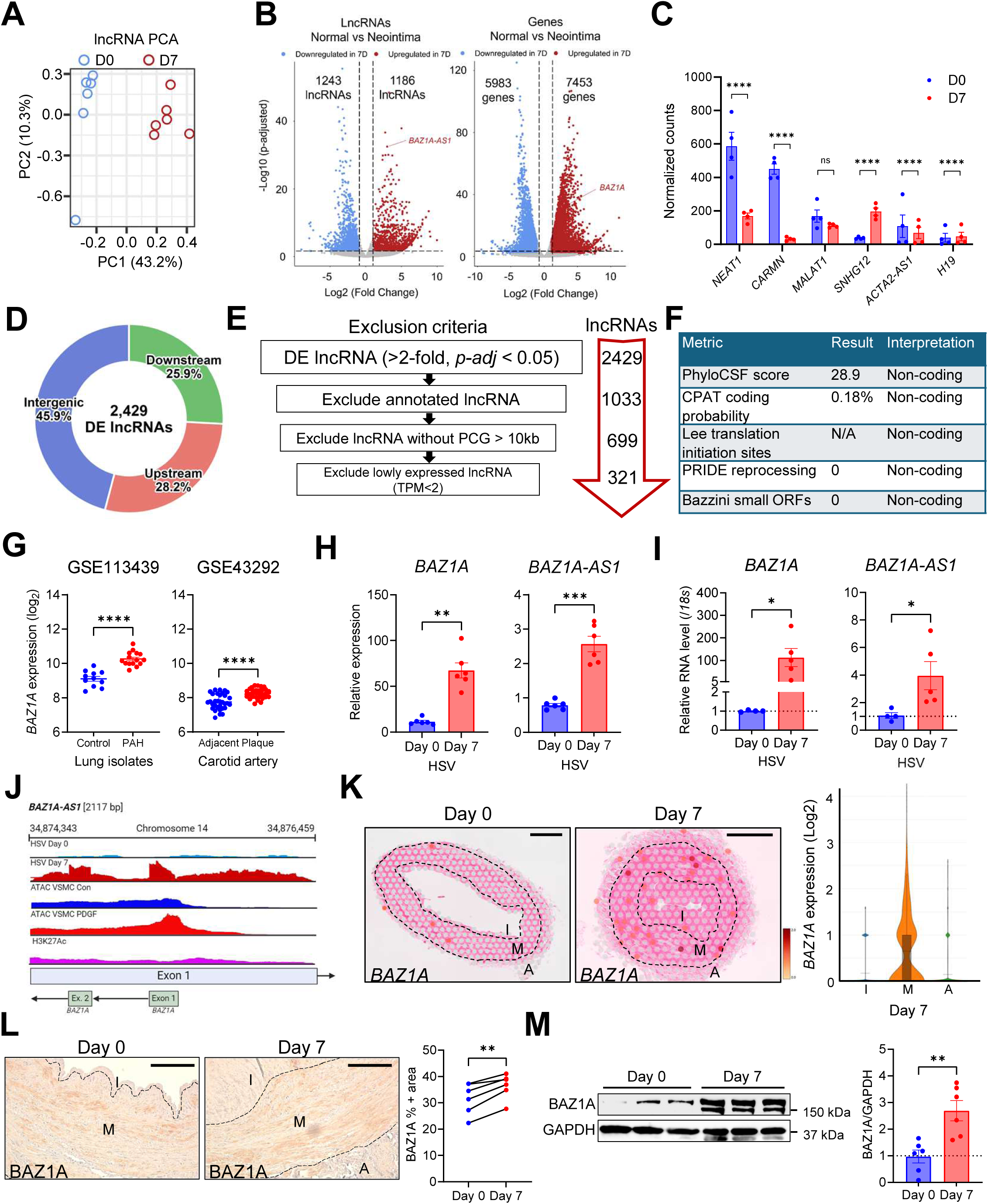
*BAZ1A-AS1*, a novel long non-coding RNA, and its *cis*-regulatory gene *BAZ1A* are upregulated in human saphenous vein during neointima proliferation. **A**, Principal component analysis of bulk RNA-seq in human saphenous vein (HSV) at Day 0 and Day 7 (n = 6). **B**, Volcano plots of differentially expressed (DE) lncRNAs (**left**) and protein-coding genes (**right**) between Day 0 and Day 7 HSV (*P adj* < 0.05, |fold change| > 2.0; n = 6). **C**, Relative expression of cardiovascular-associated lncRNAs (*NEAT1*, *CARMN*, *MALAT1*, *SNHG12*, *ACTA2-AS1*, *H19*) from RNA-seq of Day 0 vs Day 7 HSV (n = 4). **D**, Genomic distribution of DE lncRNAs relative to nearest protein-coding genes. **E**, Filtering strategy identifying 321 candidate *cis*-regulatory lncRNAs from 2,429 DE lncRNAs. **F**, Coding-potential analyses (PhyloCSF, CPAT, Lee, PRIDE, Bazzini) classify *BAZ1A-AS1* as non-coding. **G**, *BAZ1A* expression in publicly available microarrays: pulmonary arterial hypertension vs control lung (GSE113439, **left**) and carotid plaque vs adjacent tissue (GSE43292, **right**). *BAZ1A* and *BAZ1A-AS1* expression in Day 0 vs Day 7 HSV by bulk RNA-seq (**H**, n = 6) and RT-qPCR (**I**, n = 4–5). **J**, RNA-seq, ATAC-seq, and H3K27ac ChIP-seq tracks at the *BAZ1A-AS1*/*BAZ1A* locus; ATAC and ChIP data from GSE72696 (HCASMCs). **K**, Visium spatial transcriptomics feature plots of *BAZ1A* expression in Day 0 (left) and Day 7 (middle) HSV sections (scale bar: 500 μm), and quantification of *BAZ1A* expression across intima (I), media (M), and adventitia (A) (right). **L**, Representative immunohistochemistry and paired quantification of BAZ1A in Day 0 vs Day 7 HSV (n = 5). Scale bar = 200 μm. **M**, Western blot and mean densitometry of BAZ1A normalized to GAPDH in Day 0 vs Day 7 HSV (n = 6). Data presented as mean ± SEM. Statistical tests: two-tailed paired *t-*test (**I**, **L**, **M**); Wald test with Benjamini-Hochberg correction (**B**, **C**, **H**). *\*P* < 0.05, \*\**P* < 0.01, \*\*\**P* < 0.001, \*\*\*\**P* < 0.0001; ns, not significant.

Among the DE lncRNAs, 45.9% were intergenic, with 25.9% located downstream and 28.2% upstream of the nearest protein-coding gene (**Figure 1D**). To identify novel *cis*-regulatory lncRNAs that potentially regulate NP, we applied sequential filtering: exclusion of annotated lncRNAs, retention of only those with predicted *cis*-regulatory genes within 10 kb, and a minimum expression threshold of 2 transcripts per million (TPM), yielding 321 high-confidence candidates (**Figure 1E**). Among these, *AL121603.2* was amongst the most highly upregulated at Day 7. This transcript is a 2,116-nt RNA transcribed antisense to the 5′ end of the *Bromodomain Adjacent to Zinc Finger 1A* (*BAZ1A*) gene on chromosome 14 (chr14(+):34,874,343–34,876,459) and was named *BAZ1A-antisense 1* (*BAZ1A-AS1*) according to HUGO guidelines. *In silico* analysis using PhyloCSF, CPAT, Lee translation initiation site prediction, PRIDE reprocessing, and Bazzini small ORF detection uniformly classified *BAZ1A-AS1* as non-coding (**Figure 1F**).

Moreover, *BAZ1A* expression was elevated in human lung tissue from patients with pulmonary arterial hypertension compared to controls (GSE113439)^23^ and in human carotid artery plaque compared to adjacent tissue (GSE43292)^24^ (**Figure 1G**), suggesting that *BAZ1A* upregulation is a conserved feature of two different human vascular diseases. Expression of both *BAZ1A-AS1* and *BAZ1A* was significantly higher in Day 7 compared to Day 0 HSV, as confirmed by both bulk RNA-seq (**Figure 1H**) and RT-qPCR (**Figure 1I**). RNA-seq tracks at the *BAZ1A-AS1*/*BAZ1A* locus showed increased read enrichment at Day 7, and ATAC-seq of PDGF-BB–treated HCASMCs demonstrated increased chromatin accessibility at the locus, while H3K27ac ChIP-seq showed modest enrichment at the *BAZ1A* promoter region (**Figure 1J**).^25^

Visium spatial transcriptomics demonstrated that *BAZ1A* expression was strongly enriched in the tunica media of HSV at both Day 0 and Day 7, with a significant further increase at Day 7 (**Figure 1K**). Consistent with medial VSMC enrichment, immunohistochemistry confirmed increased BAZ1A protein in Day 7 compared to Day 0 HSV (**Figure 1L**), and Western blot analysis demonstrated significantly elevated BAZ1A protein normalized to GAPDH (**Figure 1M**). Together, these data identify *BAZ1A-AS1* as a novel VSMC-enriched lncRNA that is coordinately upregulated with its predicted *cis*-regulatory gene *BAZ1A* during NP in HSV.

### Characterization of *BAZ1A-AS1* and induction of the *BAZ1A-AS1*/*BAZ1A* axis by UV exposure

3′ Rapid Amplification of cDNA Ends (RACE) analysis in HSVSMCs confirmed a single 2,116-nt isoform of *BAZ1A-AS1* (**Figure 2A**). Consistent with VSMC enrichment, RNA-FISH of *BAZ1A-AS1* in Day 7 HSV showed strong expression in the tunica media and, to a lesser extent, in the proliferating neointima (**Figure 2B**). Both *BAZ1A-AS1* and *BAZ1A* were enriched in the nucleus of HSVSMCs, as observed by RNA-FISH (**Figure 2C**) and nuclear-cytoplasmic fractionation (**Figure 2D**), with *MALAT1* serving as a nuclear-localized positive control.

**Figure 2.**
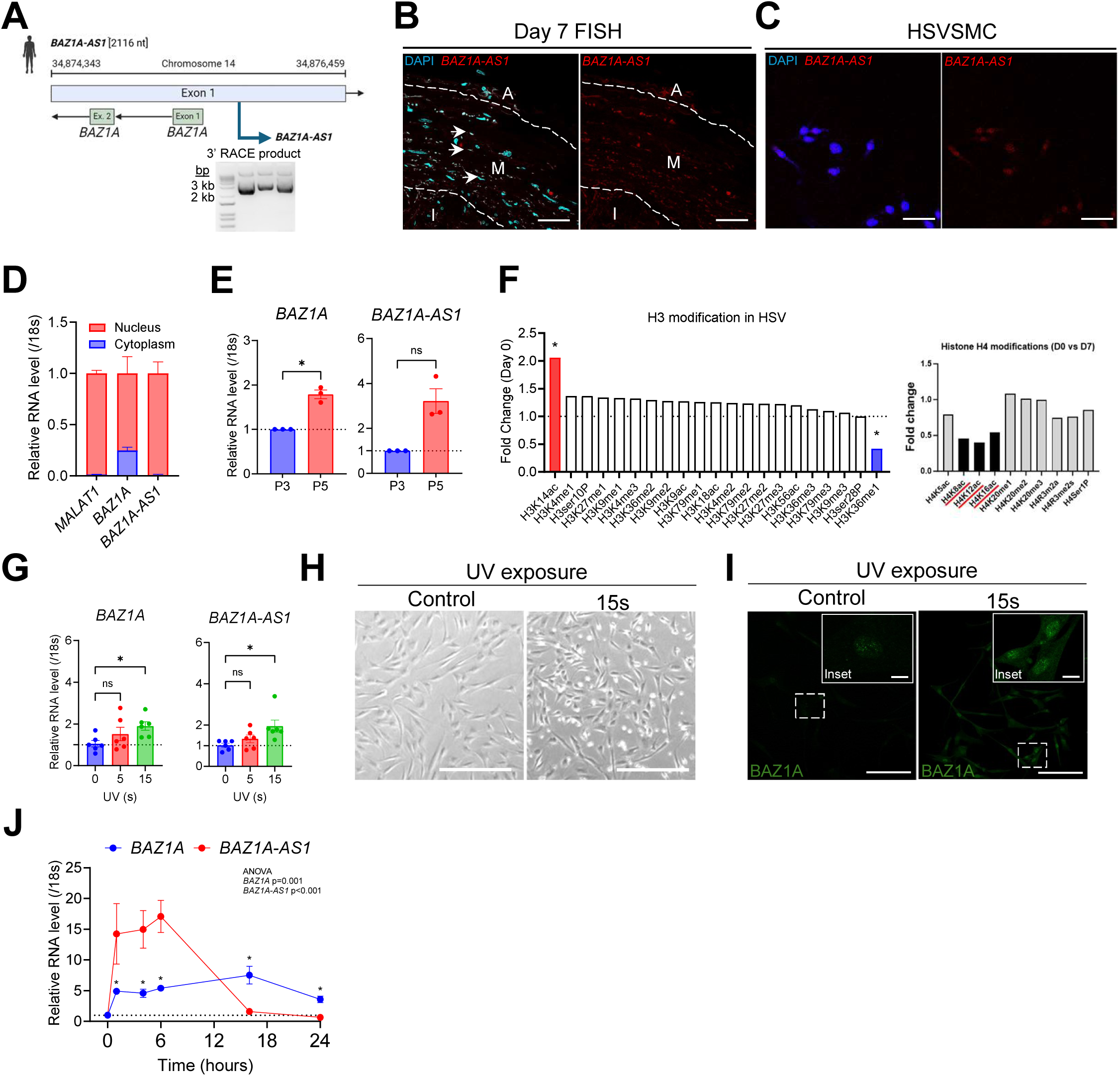
Characterization of *BAZ1A-AS1* and induction of the *BAZ1A-AS1*/*BAZ1A* axis by stimuli. **A**, Genomic locus of *BAZ1A-AS1* and *BAZ1A* on chromosome 14 with 3′ RACE product of *BAZ1A-AS1* in human saphenous vein smooth muscle cells (HSVSMCs), confirming a 2,116-nucleotide (nt) transcript. **B**, Representative RNA Stellaris FISH of *BAZ1A-AS1* in Day 7 HSV. Arrows indicate BAZ1A-AS1 signal in the tunica media. A: adventitia, M:media, I:intima. Scale bar = 100 μm. **C**, RNA FISH of *BAZ1A-AS1* in HSVSMCs. Scale bar = 50 μm. **D**, Nuclear and cytoplasmic fractionation followed by RT-qPCR for *MALAT1* (nuclear control), *BAZ1A*, and *BAZ1A-AS1* in HSVSMCs (n = 4). **E**, RT-qPCR of *BAZ1A* and *BAZ1A-AS1* in passage 3 vs passage 5 HSVSMCs (n = 3). **F**, H3 (**left**) and H4 (**right**) histone modification profiling of Day 0 vs Day 7 HSV, expressed as fold change vs Day 0 (n = 3). Red and blue bars indicate significantly altered modifications. **G**, RT-qPCR of *BAZ1A* and *BAZ1A-AS1* in HSVSMCs after 0, 5, or 15 s of UV exposure (n = 6). **H**, Representative brightfield images of HSVSMCs 24 h after control (**left**) or 15-s UV exposure (**right**). Scale bar = 100 μm. **I**, Representative immunofluorescence of BAZ1A in HSVSMCs 6 h after control (**left**) or 15-s UV exposure (**right**). Scale bar = 100 μm (main) and 10 μm (inset). **J**, Time-course RT-qPCR of *BAZ1A* and *BAZ1A-AS1* in HSVSMCs after 15-s UV exposure (n = 3). One-way ANOVA: BAZ1A *P* = 0.001; *BAZ1A-AS1 P* < 0.001. Data presented as mean ± SEM. Statistical tests: two-tailed paired *t*-test (**D**, **E**, **F**). Dunnett’s multiple-comparisons test vs 0-s control (**G**); one-way ANOVA with Welch’s *t-*test post-hoc and Benjamini-Hochberg correction (**J**).\**P* < 0.05, \*\**P* < 0.01, \*\*\*\**P* < 0.0001; ns, not significant.

To determine whether *BAZ1A-AS1* and/or *BAZ1A* expression is modulated by factors produced during NP, conditioned media (CM) collected from Day 7 HSV explants was applied to HSVSMCs. CM treatment significantly upregulated *BAZ1A* and downregulated *CNN1*, while *BAZ1A-AS1* was insignificantly changed (**Supplemental Figure 1D**). In contrast, PDGF-BB treatment significantly downregulated *BAZ1A-AS1*, with no significant change in *BAZ1A* or *CNN1* (**Supplemental Figure 1E**). Expression of *BAZ1A*, but not *BAZ1A-AS1* increased with serial passaging of HSVSMCs, with *BAZ1A* significantly elevated at passage 5 compared to passage 3 (**Figure 2E**). These findings suggest that the *BAZ1A-AS1*/*BAZ1A* axis responds to paracrine signals produced during NP in HSV explants and growth factor-induced stimulation of HSVSMCs.

To investigate epigenetic changes during NP, H3 and H4 histone modification profiling was performed on Day 0 versus Day 7 HSV. Day 7 HSV exhibited significantly increased H3K14ac and decreased H3K36me1 (**Figure 2F**). Notably, H3K14ac has been reported to be modulated by BAZ2A, a protein closely related to BAZ1A [20], and H3K36 methylation is associated with chromatin remodeling and DNA damage responses [21, 22], suggesting a potential link between NP-associated chromatin changes and BAZ1A-family proteins.

Given the established role of BAZ1A in DNA damage repair and chromatin remodeling following genotoxic stress, we tested whether UV exposure induces *BAZ1A-AS1* and *BAZ1A* expression in HSVSMCs. Both transcripts were significantly upregulated after 15 seconds of UV exposure compared to untreated controls, with no significant change at 5 seconds (**Figure 2G**). UV-exposed HSVSMCs exhibited pronounced morphological changes at 24 hours (**Figure 2H**) and increased nuclear BAZ1A protein intensity at 6 hours (**Figure 2I**). Time-course analysis following 15-s UV exposure demonstrated distinct temporal dynamics: *BAZ1A-AS1* was rapidly induced, peaking at approximately 6 hours before returning toward baseline by 18 hours, whereas *BAZ1A* expression rose more gradually and remained elevated through 24 hours (**Figure 2J**). This sequential induction pattern of transient *BAZ1A-AS1* followed by sustained *BAZ1A* is consistent with a priming role for *BAZ1A-AS1* in the early DNA damage response, preceding and potentially facilitating *BAZ1A* transcriptional activation.

### Silencing *BAZ1A-AS1* expression reduces VSMC proliferation, migration and viability

To investigate the functional role of *BAZ1A-AS1* in VSMCs, *BAZ1A-AS1* expression was silenced in HSVSMCs using locked nucleic acid (LNA) gapmeR antisense oligonucleotides (ASO), which showed efficient transfection into the nucleus (**Supplemental Figure S2C**). HSVSMCs treated with *BAZ1A-AS1* ASO (*AS1-*ASO) exhibited visibly diminished cell numbers compared to the negative control ASO (NC-ASO) (**Figure 3A**). LNA-mediated knockdown of *AS1*-ASO significantly reduced the mRNA expression of *BAZ1A-AS1* and its potential *cis*-regulatory gene *BAZ1A* under basal conditions (**Figure 3B**) and, especially, under UV exposure (**Figure 3C**).

**Figure 3.**
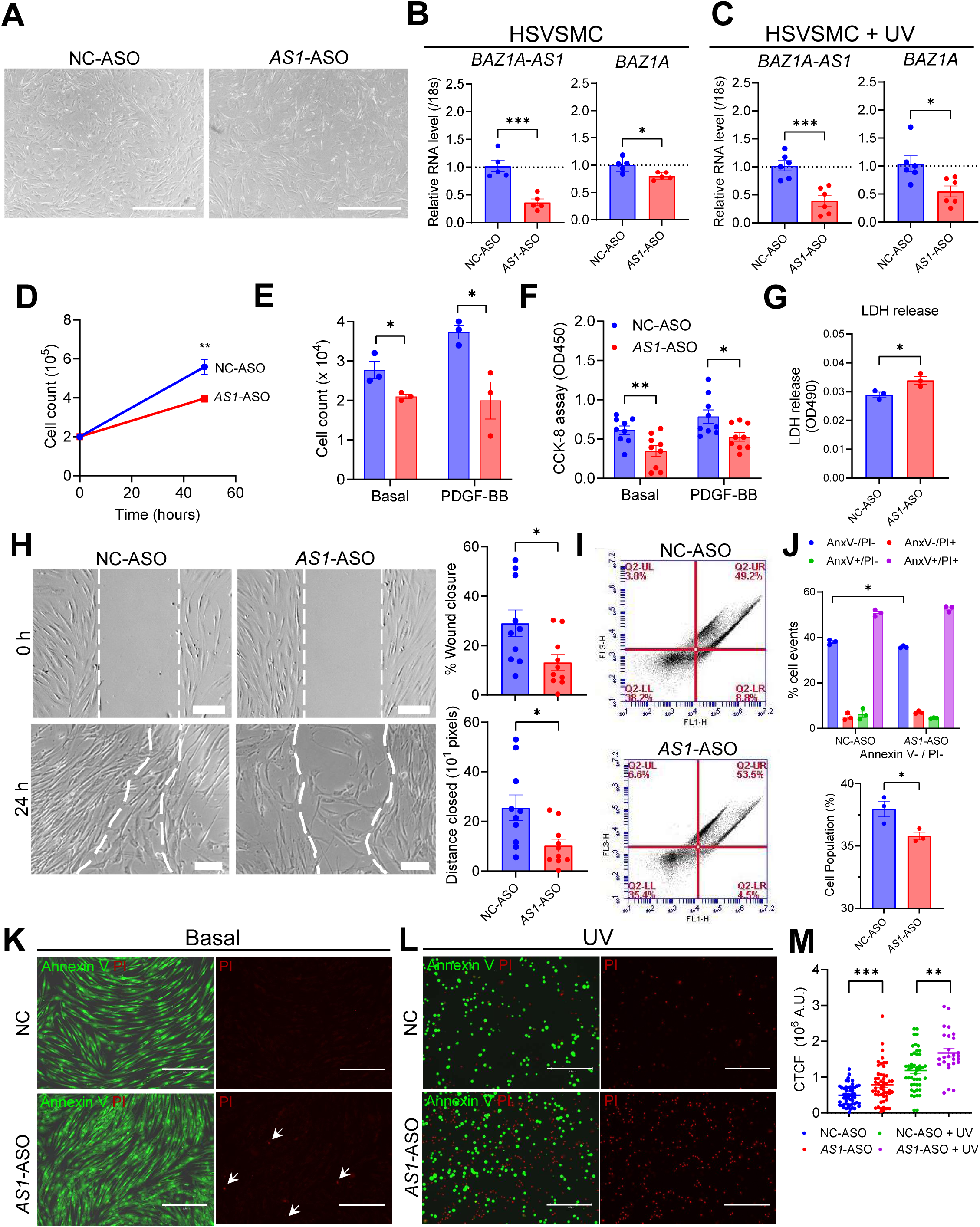
*BAZ1A-AS1* knockdown in HSVSMCs reduces proliferation, migration, and viability. **A**, Representative brightfield images of HSVSMCs 24 hours after transfection with scrambled antisense oligonucleotide (NC-ASO) or *BAZ1A-AS1* ASO. Scale bar = 200 μm. RT-qPCR of *BAZ1A-AS1* and *BAZ1A* in HSVSMCs transfected with NC-ASO or *BAZ1A-AS1* ASO under basal (**B**) and 15-s UV-exposed (**C**) conditions, harvested 24 h post-transfection (n = 5–6). **D**, Manual cell counting of HSVSMCs treated with NC-ASO or *BAZ1A-AS1* ASO (n = 5). Manual cell counting (**E**) and CCK-8 colorimetric assay (**F**) of HSVSMCs 24 h after transfection under basal and PDGF-BB (10 ng/mL) conditions (n = 3). **G**, LDH release assay 24 h after transfection (n = 3). **H**, Wound-scratch migration assay: representative brightfield images at 0 h and 24 h (left), with quantification of wound closure (%) and distance closed (right; n = 10–12). Scale bar = 30 μm. **I**, Representative Annexin V / propidium iodide (PI) flow cytometry plots 24 h after transfection. **J**, Quantification of AnxV⁻/PI⁻ (viable), AnxV⁺/PI⁻ (early apoptotic), AnxV⁻/PI⁺ (necrotic), and AnxV⁺/PI⁺ (late apoptotic) populations (top), with the AnxV⁻/PI⁻ fraction shown separately (bottom; n = 3). Representative Annexin V (green) and PI (red) immunofluorescence of HSVSMCs under basal (**K**) and 15-s UV-exposed (**L**) conditions; arrows indicate PI⁺ cells. Scale bar = 200 μm. **M**, Quantification of PI fluorescence intensity (corrected total cell fluorescence, CTCF) across NC, *BAZ1A-AS1* ASO, NC + UV, and *BAZ1A-AS1* ASO + UV groups (n = 5–6). Data presented as mean ± SEM. Statistical tests: two-tailed unpaired *t*-test (**B**, **C**, **D**, **E**, **F**, **G**, **H**, **J**); two-way ANOVA with Tukey post-hoc (**M**). \**P* < 0.05, \*\**P* < 0.01, \*\*\**P* < 0.001, \*\*\*\**P* < 0.0001; ns, not significant.

*BAZ1A-AS1* knockdown significantly reduced HSVSMC numbers as measured by manual cell counting over 48 h (**Figure 3D**). This reduction was further confirmed at 24 h under both basal and PDGF-BB–stimulated conditions by both cell counting (**Figure 3E**) and CCK-8 colorimetric assay (**Figure 3F**), with the reduction approximately 50% greater in the presence of PDGF-BB. LDH assay showed a modest but statistically significant increase in cytotoxicity in *AS1-*ASO treated HSVSMCs compared to controls (**Figure 3G**). Together, these findings suggest that knockdown of *BAZ1A-AS1* diminishes both proliferation and viability of HSVSMCs, consistent with the marked downregulation of proliferation markers *CCND1* and *CDK4* observed by RNA-seq (**Figure 6A**).

To assess migration, wound-scratch assays were performed in serum-starved HSVSMCs. Wound closure and distance closed were both significantly reduced in *BAZ1A-AS1* knockdown cells compared to NC-ASO controls (**Figure 3H**). Annexin V and propidium iodide (PI) apoptosis assays were performed by flow cytometry to evaluate cell viability and apoptosis. Knockdown of *BAZ1A-AS1* in HSVSMCs significantly reduced cell viability (AnxV⁻/PI⁻) under basal conditions, without significant differences in early or late apoptotic populations (**Figure 3I, J**).

To further characterize cell death in the context of genotoxic stress, Annexin V and PI immunofluorescence was performed on HSVSMCs under basal (**Figure 3K**) and 15-s UV-exposed (**Figure 3L**) conditions. Reduced cell density in the UV-exposed panels likely reflects a combination of UV-induced cell loss and morphological changes associated with genotoxic stress, as observed in brightfield imaging. Quantification of PI fluorescence demonstrated significantly increased PI signal in *AS1*-ASO–treated HSVSMCs, which was further augmented by UV exposure (**Figure 3M**). Collectively, these data suggest that *BAZ1A-AS1* knockdown reduces VSMC proliferation and migration while increasing susceptibility to cell death, consistent with a key role for this lncRNA in regulating multiple VSMC responses during NP.

### *BAZ1A* knockdown impacted VSMC functions in a manner that resembled BAZ1A-AS1 deficiency

Our data suggests that *BAZ1A-AS1* modulates *BAZ1A* expression through a *cis*-regulatory manner. To test whether *BAZ1A* itself impacts VSMC functions, *BAZ1A* was silenced by siRNA, rather than ASO, in HSVSMCs, as siRNA-mediated degradation targets mature cytoplasmic mRNA whereas LNA gapmeRs preferentially degrade nuclear transcripts. Brightfield imaging demonstrated visibly reduced cell density in si*BAZ1A*-treated HSVSMCs compared to siScram controls (**Figure 4A**). RT-qPCR confirmed that si*BAZ1A* reduced *BAZ1A* mRNA by approximately 90% and *BAZ1A-AS1* by approximately 50%, suggesting potential bidirectional regulation within the lncRNA-gene pair (**Figure 4B**). EdU-Amplex Red proliferation assay confirmed significantly reduced proliferation in si*BAZ1A*-treated HSVSMCs compared to siScram and non-transfected controls (**Figure 4C**). Notably, *BAZ1A* knockdown caused significant cytotoxicity in HSVSMCs (data not shown), precluding sustained downstream analyses in primary cells. We therefore used rhabdomyoblast (RD) cells, which tolerated *BAZ1A* silencing, for subsequent functional characterization. In RD cells, si*BAZ1A* significantly reduced cell numbers under both basal and UV-exposed conditions as measured by manual cell counting over 48 h (**Figure 4D**). CCK-8 viability assay confirmed reduced viability in si*BAZ1A*-treated RD cells under both basal and UV-exposed conditions (**Figure 4E**).

**Figure 4.**
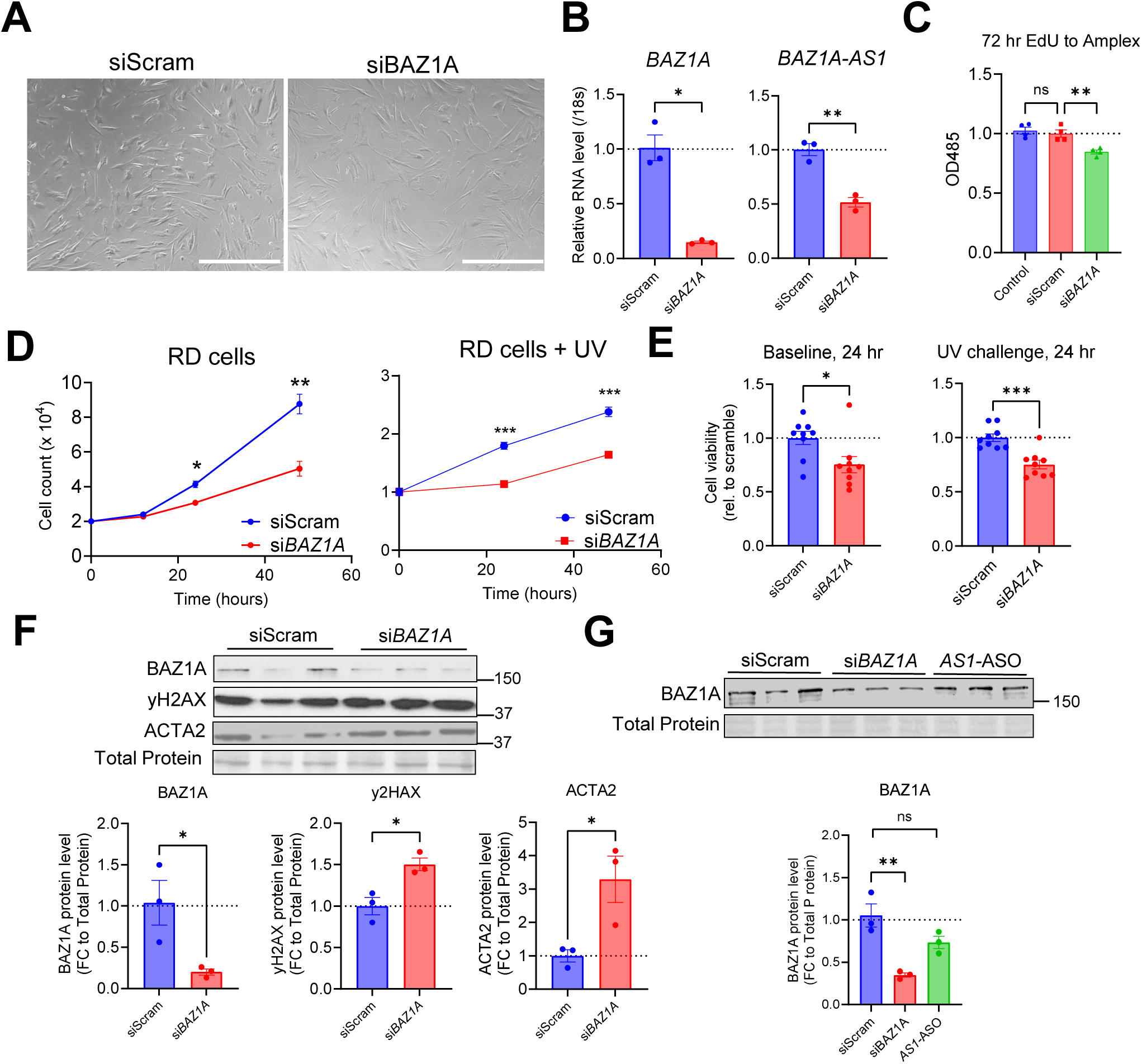
*BAZ1A* knockdown in HSVSMCs resembles *BAZ1A-AS1* knockdown to reduce cell proliferation and viability. **A**, Representative brightfield images of HSVSMCs 48 h after transfection with scrambled siRNA (siScram) or *BAZ1A* siRNA (si*BAZ1A*). Scale bar = 100 μm. **B**, RT-qPCR of *BAZ1A* and *BAZ1A-AS1* in HSVSMCs treated with siScram or si*BAZ1A* (n = 3). **C**, EdU-Amplex Red proliferation assay in HSVSMCs 72 h after transfection with control, siScram, or si*BAZ1A* (n = 3). **D**, Manual cell counting of rhabdomyoblast (RD) cells treated with siScram or si*BAZ1A* under basal (left) and 15-s UV-exposed (right) conditions (n = 4). **E**, CCK-8 colorimetric assay of RD cells 24 h after transfection under basal (left) and 15-s UV-exposed (right) conditions (n = 4–5). **F**, Western blot (**top**) and mean densitometry analysis (**bottom**) of BAZ1A, γH2AX, and ACTA2 in HSVSMCs treated with siScram or si*BAZ1A*, normalized to total protein (n = 3). **G**, Western blot (**top**) and mean densitometry analysis (**bottom**) of BAZ1A in HSVSMCs 48 h after transfection with siScram, si*BAZ1A*, or *AS1*-ASO, normalized to total protein (n = 3). Data presented as mean ± SEM. Statistical tests: two-tailed unpaired *t*-test (**B**, **E**, **F**); one-way ANOVA with Dunnett’s multiple comparison test (**C, G**); two-way ANOVA with Tukey post-hoc (**D**). \**P* < 0.05, \*\**P* < 0.01, \*\*\**P* < 0.001, \*\*\*\**P* < 0.0001; ns, not significant.

At the protein level, Western blot analysis of HSVSMCs harvested 48 h after si*BAZ1A* transfection, prior to the onset of severe cytotoxicity, confirmed effective BAZ1A protein reduction and demonstrated significantly increased γH2AX, a DNA damage response protein, alongside significantly increased ACTA2, a marker of VSMC contractile identity (**Figure 4F**). The upregulation of ACTA2 following *BAZ1A* silencing is consistent with a role for BAZ1A in driving VSMC dedifferentiation, and its loss partially restoring the contractile phenotype, while increased γH2AX suggests accumulation of unrepaired DNA damage in the absence of BAZ1A, consistent with its established role in chromatin remodeling and nucleotide excision repair.

To determine whether *BAZ1A-AS1* knockdown affects BAZ1A protein expression, we performed Western blot analysis in HSVSMCs. Si*BAZ1A* produced robust reduction in BAZ1A protein, whereas *AS1*-ASO caused a modest, non-significant trend toward reduced BAZ1A protein (**Figure 4G**). This finding is consistent with the partial *BAZ1A* mRNA reduction observed after *BAZ1A-AS1* knockdown (**Figure 3B, C**), suggesting that *BAZ1A-AS1* may regulate VSMC function through mechanisms beyond BAZ1A protein modulation alone.

### *BAZ1A-AS1* and *BAZ1A* knockdown transcriptomes converge on suppression of proliferative and inflammatory programs

To address the potential underlying mechanisms by which *BAZ1A-AS1* regulates VSMC function, we performed bulk RNA-seq of HSVSMCs treated with NC-ASO (**Figure 5**) or *AS1*-ASO and of RD cells treated with siScram or si*BAZ1A* under UV exposure (**Supplemental Figure S3**). Principal component analysis (PCA) confirmed strong separation of knockdown from control transcriptomes (**Figure 5A**). In HSVSMCs with *BAZ1A-AS1* knockdown, 1,291 genes were upregulated and 1,191 genes were downregulated (*P-*adj < 0.05, |log2 fold change| > 0.585) **(Figure 5B, left panel**). *BAZ1A* knockdown in RD cells produced a more focused transcriptional response, with 592 upregulated and 637 downregulated genes, accompanied by significant downregulation of both *BAZ1A* and *BAZ1A-AS1* **(Figure 5B, right panel**). Heatmap analysis of NC-ASO versus *AS1*-ASO HSVSMCs demonstrated coordinated transcriptional shifts across five functional programs: inflammatory signaling (*IL6, CXCL1*), cell cycle control (*CCND1, CDK1, E2F2, MKI67*), stress response (*DDIT3, CHAC1, TRIB3, SESN2*), VSMC differentiation (*CNN1, ACTA2, LMOD1, SYNPO2*) and extracellular matrix remodeling (*MMP2, MMP14, CTNNB1*) (**Figure 5C**).

**Figure 5.**
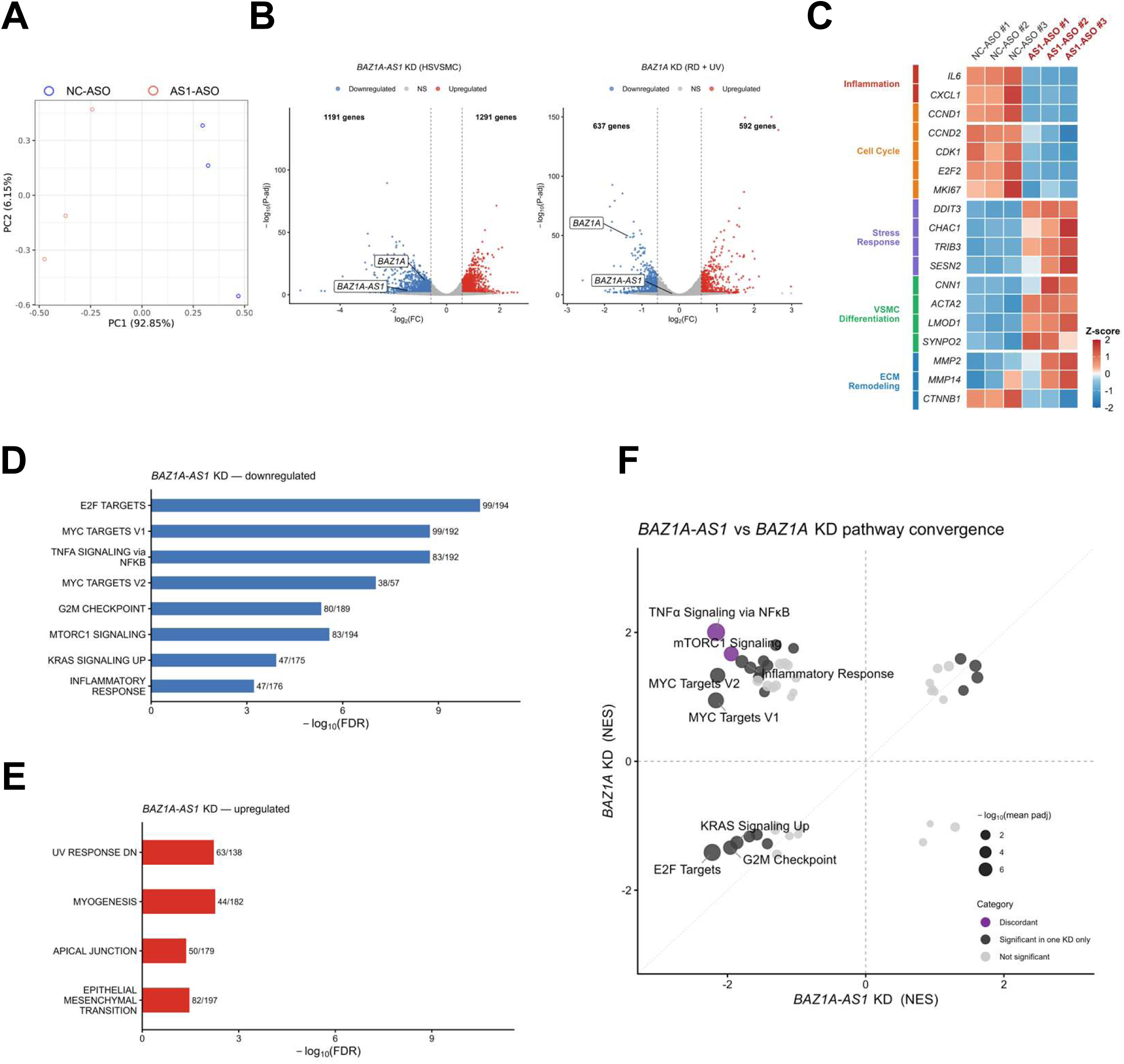
*BAZ1A-AS1* and *BAZ1A* knockdown transcriptomes converge on suppression of proliferative and inflammatory programs in VSMCs. **A**, Principal component analysis of bulk RNA-seq in HSVSMCs transfected with NC-ASO or *BAZ1A-AS1* ASO (*AS1*-ASO) (n = 3 per group). **B**, Volcano plots of DE genes in HSVSMCs treated with *AS1*-ASO (**left**) and in RD cells treated with siScram or si*BAZ1A* under UV exposure (**right**), with thresholds *P-*adj < 0.05 and |fold change| > 0.585. **C**, Heatmap of z-scored expression of representative genes across five functional categories (Inflammation, Cell Cycle, Stress Response, VSMC Differentiation, ECM Remodeling) in NC-ASO versus *AS1*-ASO HSVSMCs (n = 3 per group). Hallmark gene set enrichment analysis (GSEA) of *BAZ1A-AS1* knockdown transcriptome showing top downregulated (**D**) and upregulated (**E**) pathways, ranked by −log10(FDR). Bar labels indicate the number of leading-edge genes over total pathway gene set size. **F**, Pathway-level convergence between *BAZ1A-AS1* and *BAZ1A* knockdown transcriptomes. Each point represents one Hallmark gene set, plotted by normalized enrichment score (NES) in *-AS1* KD (HSVSMCs, x-axis) versus *BAZ1A* KD (RD cells under UV, y-axis). Purple points indicate pathways with significant enrichment (FDR < 0.05) and discordant direction; dark grey points indicate pathways significant in one condition only; light grey points indicate pathways not significant in either. Point size reflects combined significance (−log10 of the geometric mean of adjusted p-values). Dashed lines indicate zero NES; dotted line indicates y = x. Statistical tests: Wald test with Benjamini-Hochberg correction (**B**); pre-ranked GSEA with 1,000 permutations and Benjamini-Hochberg FDR correction (**D, E, F**).

Hallmark gene set enrichment analysis (GSEA) of the *BAZ1A-AS1* knockdown transcriptome identified strong downregulation of proliferative pathways (E2F targets, MYC targets, G2M checkpoint), inflammatory pathways (TNFα signaling via NFκB, Inflammatory Response), and stress-response programs (mTORC1 signaling, KRAS signaling) (**Figure 5D**). Upregulated pathways were more modest, including myogenesis, Epithelial-Mesenchymal Transition, and UV response DN (**Figure 5E**).

To evaluate whether the knockdown of *BAZ1A-AS1* and *BAZ1A* converge on a shared transcriptional program, we plotted normalized enrichment scores (NES) for each Hallmark pathway across the two knockdown contexts (**Figure 5F**). Most proliferative and inflammatory pathways were concordantly suppressed, clustering linearly along the diagonal. Two pathways, TNFα signaling via NFκB and mTORC1 signaling, showed discordant direction between knockdown of *BAZ1A-AS1* and *BAZ1A*. These findings suggest that, for the most part, *BAZ1A-AS1* and *BAZ1A* function within a common regulatory axis. Knockdown of either transcript suppressed proliferative and stress-response transcriptional programs in HSVSMCs (*BAZ1A-AS1* KD) and RD cells (*BAZ1A* KD), indicating convergent downstream effects across both cell types.

### BAZ1A-AS1 physically interacts with the BAZ1A locus and selectively engages phenotype-relevant transcripts

To validate key transcriptomic findings, RT-qPCR was performed in HSVSMCs with *BAZ1A-AS1* or *BAZ1A* knockdown. *BAZ1A-AS1* knockdown significantly increased expression of VSMC differentiation markers *CNN1* and *TAGLN*, while reducing proliferation markers *CCND1* and *CDK4* (**Figure 6A**). *BAZ1A* knockdown also significantly increased *CNN1* expression, with a trend towards increased *TAGLN* (p=0.08), and significantly reduced *CCND1,* while additionally upregulating *ACTA2* (**Figure 6B**). The concordant direction of these changes across both knockdowns further supports a shared regulatory axis between *BAZ1A-AS1* and *BAZ1A* in controlling VSMC phenotype.

**Figure 6.**
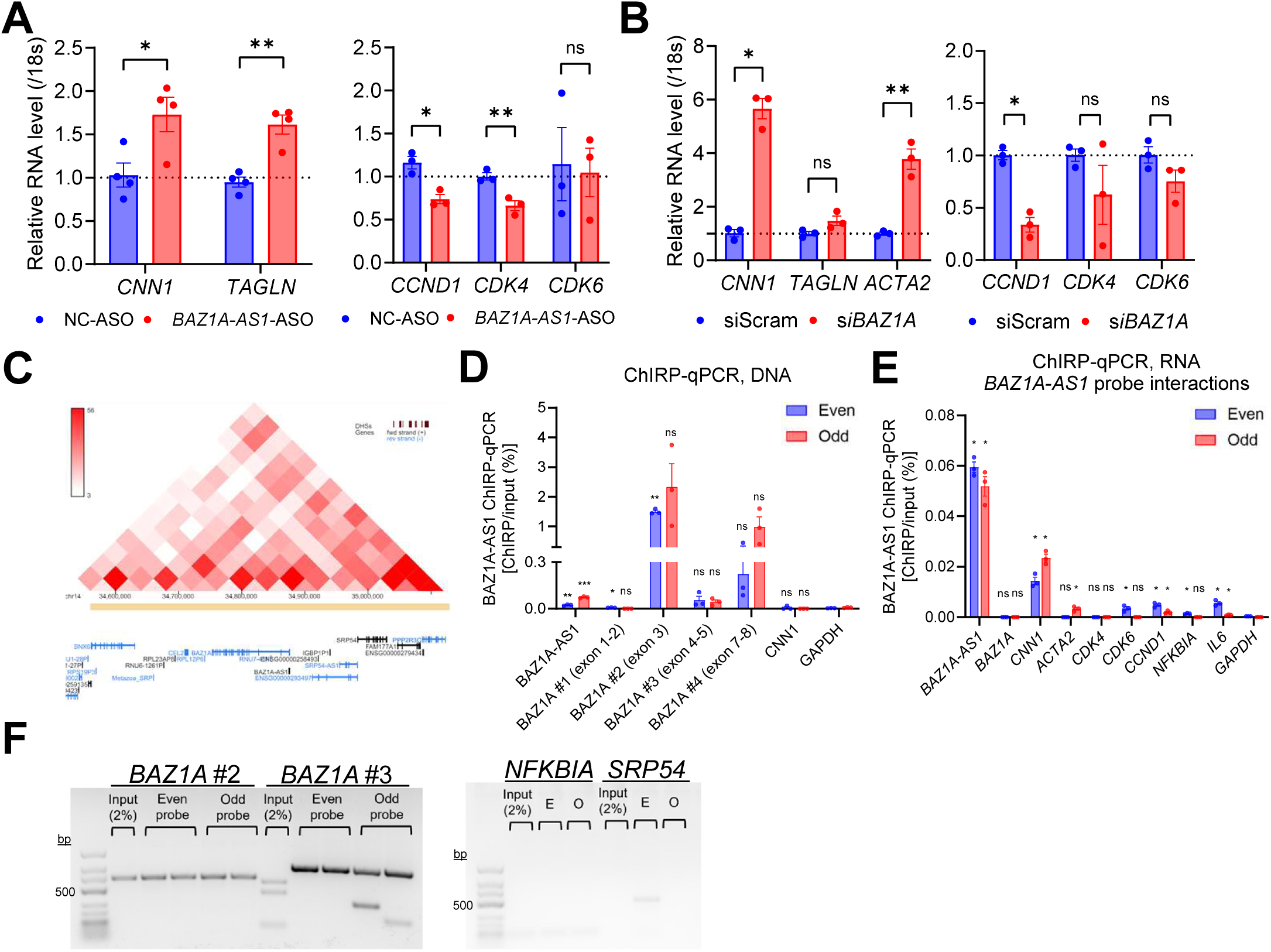
*BAZ1A-AS1* physically interacts with the *BAZ1A* locus and selectively engages phenotype-relevant transcripts in VSMCs. **A**, RT-qPCR validation of VSMC differentiation (*CNN1*, *TAGLN*) and proliferation (*CCND1*, *CDK4*, *CDK6*) markers in HSVSMCs transfected with NC-ASO or *BAZ1A-AS1* ASO (*AS1*-ASO) (n = 3–4). **B**, RT-qPCR of VSMC differentiation (*ACTA2*, *CNN1*, *TAGLN*) and proliferation (*CCND1*, *CDK4*, *CDK6*) markers in HSVSMCs transfected with siScram or si*BAZ1A* (n = 3). **C**, Hi-C contact map of the human aorta genome across a 500-kb window centered on the *BAZ1A-AS1*/*BAZ1A* locus (chr14), with the RefSeq gene track shown below; Hi-C data obtained from GEO: GSE58752. **D**, ChIRP-qPCR of DNA pulldown in HCASMCs using *BAZ1A-AS1* probes split into Even and Odd pools, targeting *BAZ1A-AS1* (bait control), *BAZ1A* exons 1–2, 3, 4–5, and 7–8, *CNN1*, and *GAPDH* (negative control); enrichment expressed as percent input (n = 3 per pool). **E**, ChIRP-qPCR of RNA pulldown in HCASMCs using Even and Odd *BAZ1A-AS1* probe pools against a panel of candidate interacting transcripts (*BAZ1A*, *CNN1*, *ACTA2*, *CDK4*, *CDK6*, *CCND1*, *NFKBIA*, *IL6*) with *GAPDH* as negative control (n = 3 per pool). **F**, Representative gel of ChIRP-qPCR DNA products (28 cycles) for *BAZ1A* exon 3 (#2), exon 4–5 (#3), *NFKBIA*, and *SRP54* showing Even (E) and Odd (O) probe pool amplification relative to 2% input. Data presented as mean ± SEM. Statistical tests: two-tailed unpaired **t**-test (**A**, **B**); Welch’s unpaired one-sided *t*-test (target > GAPDH) with Benjamini-Hochberg correction across all target–pool comparisons (**D**, **E**). \**P* < 0.05, \*\**P* < 0.01, \*\*\**P* < 0.001, \*\*\*\**P* < 0.0001; ns, not significant.

To investigate the physical basis of this regulation, we examined Hi-C contact data from human aorta (GEO: GSE58752)^16^ within a 500-kilobases window centered on the *BAZ1A-AS1*/*BAZ1A* locus. The contact map demonstrated frequent chromatin interactions between *BAZ1A-AS1* and several neighboring genes, including *BAZ1A, NFKBIA,* and *SRP54* (**Figure 6C**). To determine whether *BAZ1A-AS1* physically interacts with these loci, we performed ChIRP-qPCR using biotinylated antisense probes targeting *BAZ1A-AS1*, split into Even and Odd pools as internal controls. Because *BAZ1A-AS1* is transcribed antisense to the 5′ end of *BAZ1A* exon 1, probes were designed to exclude this overlapping region to avoid artifactual signal from direct antisense complementarity. As expected, DNA pulldown showed low enrichment at *BAZ1A* exons 1–2, the region closest to and partially overlapping with the *BAZ1A-AS1* locus, where probe exclusion limits detection of proximity-based interactions (**Figure 6D**). Importantly, enrichment was detected at the non-overlapping exon 3 region of *BAZ1A*, with significant pulldown in the Even probe pool, demonstrating that *BAZ1A-AS1* physically contacts downstream regions of the *BAZ1A* gene beyond its immediate site of transcription (**Figure 6D**). Significant enrichment was not detected at *NFKBIA*, and gel electrophoresis of ChIRP-qPCR products (28 cycles) confirmed specific amplification at *BAZ1A* exons 3 and 4–5 with no detectable product for *NFKBIA* or *SRP54* (**Figure 6F**).

To assess whether *BAZ1A-AS1* also engages specific mRNA transcripts, RNA obtained from ChIRP pulldown was analyzed by qPCR. *BAZ1A-AS1* selectively interacted with a subset of phenotype-relevant transcripts: *CNN1*, *CCND1*, and *IL6* mRNAs showed significant enrichment in both Even and Odd probe pools compared to *GAPDH* (**Figure 6E**). In contrast, *BAZ1A*, *CDK4*, or *GAPDH* mRNA were not significantly enriched, while *ACTA2*, *CDK6*, and *NFKBIA* were significant in only one probe pool and were therefore considered inconclusive. Notably, whereas DNA-level interactions were concentrated at the *BAZ1A* locus (**Figure 6D**), the RNA-level interactions involved a distinct and selective set of transcripts, suggesting that *BAZ1A-AS1* engages chromatin and mRNA targets through separate mechanisms.

Taken together, these findings demonstrate that *BAZ1A-AS1* physically associates with the *BAZ1A* genomic locus at non-overlapping distal exons and selectively engages a subset of phenotype-relevant transcripts, supporting a multi-level regulatory mechanism in which *BAZ1A*-*AS1* acts both through chromatin-level modulation of *BAZ1A* and through direct interactions with key mRNAs governing VSMC function.

### *Baz1a* is upregulated following vascular injury, and its deficiency attenuates neointima formation in mice

Data presented above were obtained from human tissues *ex vivo* and VSMCs *in vitro*. Given that *BAZ1A-AS1* is a human-specific lncRNA with no orthologue in mice, we examined the role of *Baz1a*, its protein-coding partner, in vascular injury *in vivo*. Immunofluorescence confirmed Baz1a expression in wild-type mouse aorta, co-localizing with αSma-positive VSMCs (**Figure 7A**). Analysis of a publicly available microarray dataset (GSE40637) demonstrated significant upregulation of *Baz1a* in mouse carotid arteries following wire injury (**Figure 7B**). Consistently, RT-qPCR of carotid arteries from our own complete ligation model confirmed significant *Baz1a* upregulation in ligated versus sham carotid arteries (**Figure 7C**), and immunofluorescence demonstrated increased Baz1a intensity in ligated vessels (**Figure 7D, E**). Western blot further confirmed significantly elevated Baz1a protein in ligated carotid arteries (**Figure 7F**). Collectively, these data establish that *Baz1a* is an injury-inducible gene in the mouse vasculature, paralleling the upregulation of *BAZ1A* and *BAZ1A-AS1* observed during neointima formation in human SV.

**Figure 7.**
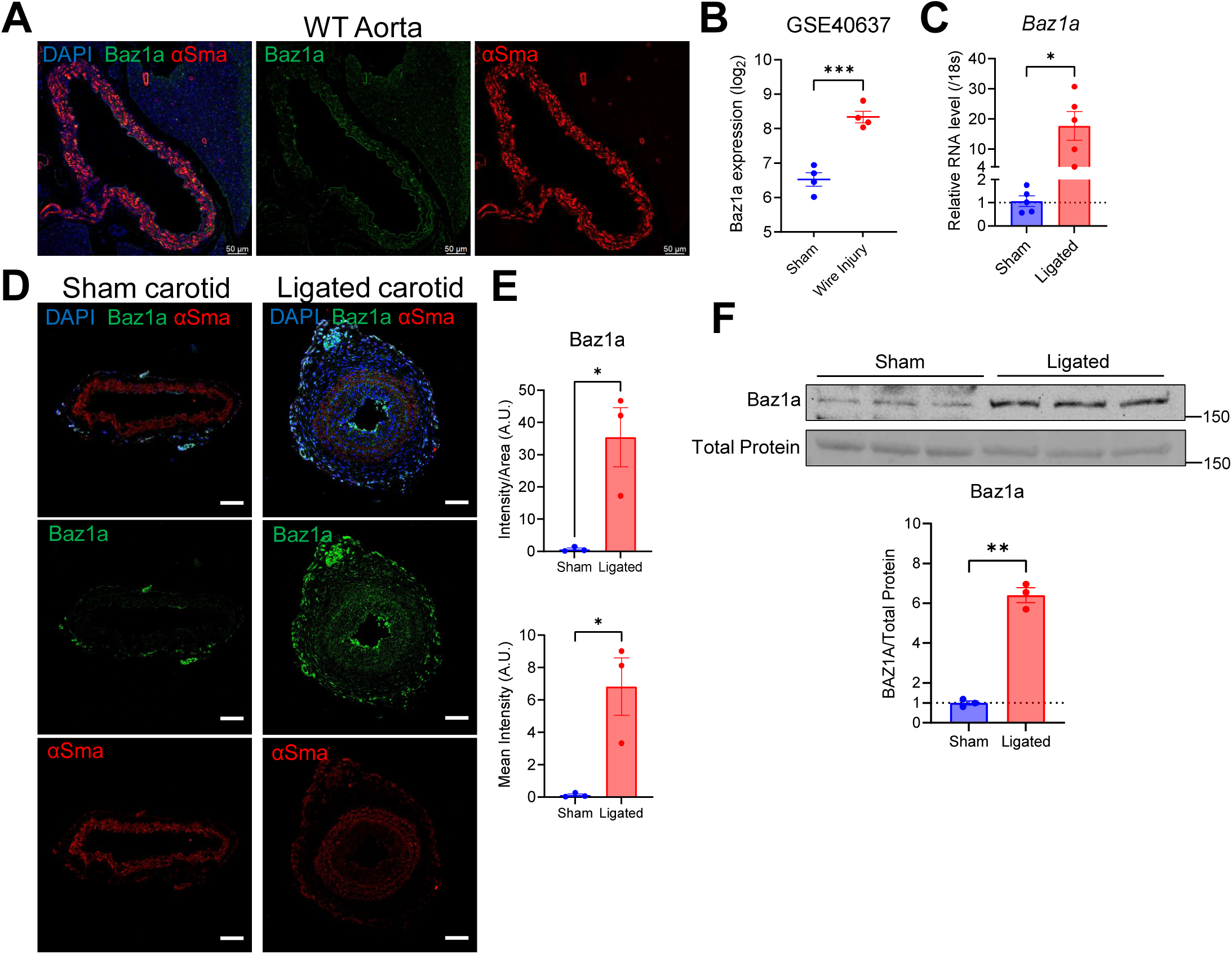
*Baz1a* is upregulated following vascular injury in mice. **A**, Representative immunofluorescence of wild-type mouse aortas co-stained for DAPI (blue), Baz1a (green), and αSma (red). Scale bar = 50 μm. **B**, *Baz1a* expression in mouse carotid arteries 14 days after sham or wire injury surgery, from a publicly available microarray dataset (GSE40637). **C**, RT-qPCR of *Baz1a* in sham and ligated carotid arteries of wild-type mice 21 days post-ligation (n = 5). **D**, Representative immunofluorescence of sham and ligated carotid arteries co-stained for DAPI (blue), Baz1a (green), and αSma (red). Scale bar = 50 μm. **E**, Quantification of Baz1a immunofluorescence intensity normalized to vessel area (top) and as mean intensity (bottom) in sham vs ligated carotid arteries (n = 3). **F**, Western blot (top) and mean densitometry analysis (bottom) of Baz1a in sham and ligated carotid arteries, normalized to total protein (n = 3). Data presented as mean ± SEM. Statistical tests: two-tailed unpaired *t*-test (**B**, **C**, **E**, **F**). *P < 0.05, \*\**P* < 0.01, \*\*\**P* < 0.001, \*\*\*\**P* < 0.0001; ns, not significant.

To determine whether Baz1a is functionally required for neointima formation, we utilized global *Baz1a* heterozygous knockout mice (*Baz1a*⁺/⁻) bred into an *Ldlr*⁻/⁻ background; homozygous knockouts were not used due to their established infertility phenotype. RT-qPCR confirmed reduced *Baz1a* expression in both aortas and left carotid arteries (LCAs) of *Baz1a*⁺/⁻;*Ldlr*⁻/⁻ mice compared to *Ldlr*⁻/⁻ controls (**Figure 8A**). Complete ligation of the left carotid artery was performed to induce neointima formation, and representative H&E sections at 200 μm and 400 μm proximal to the ligation site demonstrated markedly attenuated neointima in *Baz1a*⁺/⁻;*Ldlr*⁻/⁻ mice (**Figure 8B**). Quantification confirmed significant reductions in both neointima area and neointima-to-media ratio (**Figure 8C**), with modestly increased medial area at 400 µm but not 200 µm (**Figure 8D**).

**Figure 8.**
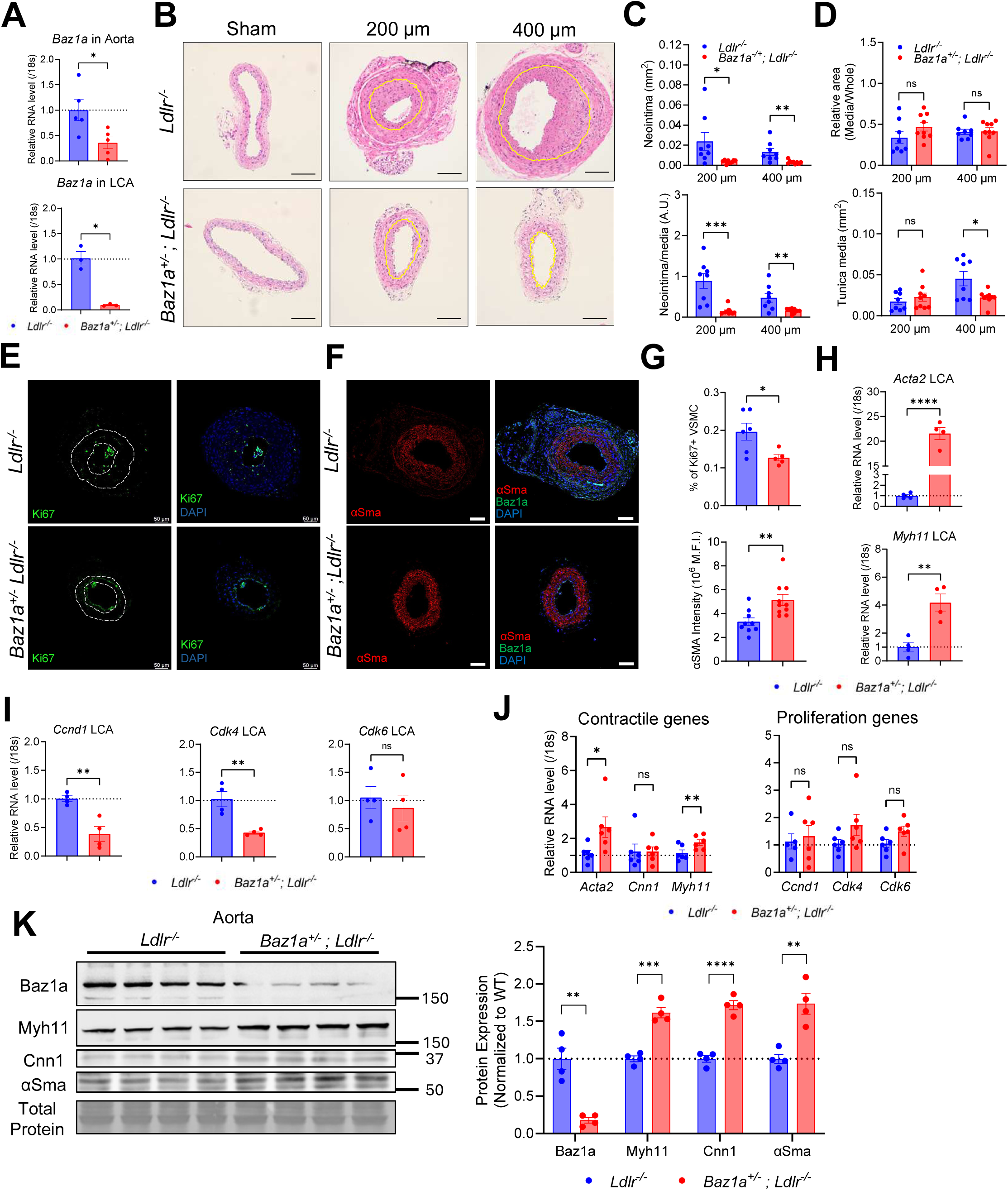
*Baz1a* heterozygosity attenuates neointima formation after carotid ligation and preserves VSMC contractile gene expression. **A**, RT-qPCR of *Baz1a* in aortas (**top**) and left carotid arteries (LCA, **bottom**) of *Ldlr*⁻fIJ⁻ and *Baz1a*⁺fIJ⁻;*Ldlr*⁻fIJ⁻ mice (n = 4, 2 mice pooled per biological replicate). **B**, Representative H&E images of sham (**left**) and ligated carotid arteries 200 μm (**middle**) and 400 μm (**right**) proximal to the ligation site, 4 weeks after complete left carotid ligation in *Ldlr*⁻fIJ⁻ and *Baz1a*⁺fIJ⁻;*Ldlr*⁻fIJ⁻ mice. Yellow dashed lines delineate the internal elastic lamina, separating the neointima (luminal side) from the tunica media. Scale bar = 100 μm. **C**, Quantification of neointima area (**top**) and neointima-to-media (N/M) ratio (**bottom**) at 200 μm and 400 μm proximal to the ligation site (n = 8–9 mice). **D**, Quantification of media/vessel area ratio (**top**) and total media area (**bottom**) at 200 μm and 400 μm proximal to the ligation site (n = 8–9 mice). **E**, Representative immunofluorescence of Ki67 (green) and DAPI (blue) in ligated carotid arteries of *Ldlr*⁻fIJ⁻ and *Baz1a*⁺fIJ⁻;*Ldlr*⁻fIJ⁻ mice. White dashed lines delineate the medial layer used for quantification of Ki67⁺ VSMCs. Scale bar = 50 μm. **F**, Representative immunofluorescence of αSma (red), Baz1a (green), and DAPI (blue) in ligated carotid arteries. Scale bar = 50 μm. **G**, Quantification of Ki67⁺ VSMCs (**top**) and αSma fluorescence intensity (**bottom**) (n = 6–8 mice). **H**, RT-qPCR of VSMC contractile genes *Acta2* and *Myh11* in ligated LCAs (n = 4, 2 mice pooled per biological replicate). **I**, RT-qPCR of proliferation genes *Ccnd1*, *Cdk4*, and *Cdk6* in ligated LCAs (n = 4, 2 mice pooled per biological replicate). **J**, RT-qPCR of contractile genes (*Acta2*, *Cnn1*, *Myh11*, left) and proliferation genes (*Ccnd1*, *Cdk4*, *Cdk6*, right) in aortas of *Ldlr*⁻/⁻ and *Baz1a*⁺/⁻;*Ldlr*⁻/⁻ mice (n = 4, 2 mice pooled per biological replicate). **K**, Western blot (**left**) and mean densitometry (**right**) of Baz1a, Myh11, Cnn1, and αSma in aortas of *Ldlr*⁻/⁻ and *Baz1a*⁺/⁻;*Ldlr*⁻/⁻ mice, normalized to total protein (n = 4 individual mice). Data presented as mean ± SEM. Statistical tests: two-tailed unpaired t-test (**A**, **C**, **D**, **G**, **H**, **I**, **J**, **K**). \**P* < 0.05, \*\**P* < 0.01, \*\*\**P* < 0.001, \*\*\*\**P* < 0.0001; ns, not significant.

To assess whether *Baz1a* deficiency modulates VSMC phenotype within the injured vessel, immunofluorescence was performed on ligated carotid arteries. The proportion of Ki67⁺ proliferating VSMCs was significantly reduced in *Baz1a*⁺/⁻;*Ldlr*⁻/⁻ mice, while αSma intensity was significantly increased (**Figure 8E–G**), indicating preserved contractile identity and reduced proliferative activity.

RT-qPCR analysis of ligated LCAs demonstrated that *Baz1a*⁺/⁻;*Ldlr*⁻/⁻ mice showed significant upregulation of VSMC contractile genes *Acta2* and *Myh11* (**Figure 8H**) and significant downregulation of proliferation genes *Ccnd1* and *Cdk4*, with *Cdk6* unchanged (**Figure 8I**), compared with *Ldlr*⁻/⁻ controls. Extended histological analysis, including elastin and collagen quantification, is shown in **Supplemental Figure S4A-E**; notably, elastin breaks were significantly increased in *Baz1a*⁺/⁻;*Ldlr*⁻/⁻ mice at both 200 μm and 400 μm, with significantly reduced elastin area fraction at 200 μm, while collagen content was unchanged, suggesting without altered extracellular matrix composition. In uninjured contralateral RCAs, *Ccnd1* and *Cdk4* were paradoxically increased despite unchanged contractile gene expression (**Supplemental Figure S4F**). Given the low baseline proliferative activity in quiescent vessels and the use of a global knockout model, these changes may reflect contributions from non-VSMC cell types and their functional significance remains unclear.

Gene expression analysis of aortas from *Baz1a*⁺/⁻;*Ldlr*⁻/⁻ mice showed significant upregulation of contractile markers *Acta2* and *Myh11* compared to control *Ldlr*⁻/⁻ mice, while *Cnn1* and proliferation markers (*Ccnd1*, *Cdk4*, *Cdk6*) were unchanged (**Figure 8J**). At the protein level, Western blot of aortic tissue confirmed significantly reduced Baz1a and increased Myh11, Cnn1, and αSma in *Baz1a*⁺/⁻;*Ldlr*⁻/⁻ mice (**Figure 8K**), providing protein-level validation of preserved contractile identity. These results demonstrate that Baz1a deficiency protects against injury-induced neointima formation in mice while preserving VSMC contractile gene expression, consistent with role for the *BAZ1A*-*AS1*/*BAZ1A* axis in VSMC phenotype and vascular remodeling.

## Discussion

This study identifies *BAZ1A-AS1*, a previously uncharacterized human-specific lncRNA, and its cis-regulatory partner *BAZ1A* as novel regulators of VSMC phenotypic modulation and NP. Four convergent lines of evidence support this conclusion. First, *BAZ1A-AS1* and *BAZ1A* were coordinately upregulated during *ex vivo* NP in HSV, with expression predominantly localized to medial and neointimal VSMCs. Second, silencing either transcript attenuated VSMC proliferation and migration while upregulating contractile markers, with concordant transcriptomic changes observed across both knockdowns. Third, ChIRP-qPCR demonstrated that *BAZ1A-AS1* engages the *BAZ1A* genomic locus at the DNA level and selectively interacts with a distinct subset of phenotype-relevant mRNAs, indicating dual regulatory modalities. Fourth, even partial reduction of *Baz1a* in heterozygous knockout mice was sufficient to attenuate neointima formation and preserve VSMC contractile identity following carotid artery ligation. Together, these findings establish the *BAZ1A-AS1/BAZ1A* axis as a stress-responsive regulatory module that promotes VSMC dedifferentiation and maladaptive vascular remodeling.

Among the 2,429 lncRNAs DE during NP in our *ex vivo* model, we prioritized *BAZ1A-AS1* based on several complementary criteria: robust coordinated upregulation with its putative cis-regulatory gene *BAZ1A* at Day 7; relatively high basal expression in HSVSMCs (∼20 copies per cell, CT ∼30); predominant localization to the VSMC-rich medial and neointimal compartments by both spatial transcriptomics and RNA-FISH; and concurrent epigenetic chromatin modifications, specifically, increased H3K14ac and decreased H3K36me1 during NP, implicating BAZ-family chromatin remodelers in this process, as H3K14ac is modulated by BAZ2A^26^ and H3K36 methylation is linked to chromatin remodeling and DNA damage responses.^27,28^ The concurrent differential expression of lncRNAs with established roles in VSMC biology, including *NEAT1,*^29^ *CARMN*,^7,8^ and *SENCR*,^30^ reinforces VSMC phenotypic switching as a central driver of NP in HSV and positions *BAZ1A-AS1* within a broader lncRNA-mediated regulatory landscape governing VSMC plasticity.

A distinguishing feature of the *BAZ1A-AS1/BAZ1A* axis is its activation by genotoxic stress rather than canonical growth factor signaling. UV exposure rapidly and robustly induced both transcripts in HSVSMCs, whereas PDGF-BB, TGFβ, TNFα, and FBS did not produce significant transcriptional upregulation. The basal expression and passage-dependent increase of *BAZ1A* in cultured HSVSMCs may reflect cumulative cellular stress inherent to *in* vitro culture conditions, including mechanical disruption during passaging and oxidative stress, rather than growth factor signaling *per se.* This stimulus-specificity aligns with the established function of BAZ1A as a cofactor of the ACF chromatin remodeling complex, which facilitates nucleotide excision repair at UV-damaged sites in an MLL1-dependent manner^31^ and partners with SNF2H (SMARCA5) to reposition nucleosomes during the DNA damage response.^31,32^ Consistent with this, *BAZ1A* silencing in HSVSMCs increased γH2AX levels, indicating accumulation of unrepaired DNA damage in the absence of BAZ1A and confirming its functional role in maintaining genomic integrity in VSMCs. The temporal dynamics of induction further support a sequential activation model: *BAZ1A-AS1* expression peaked within 6 hours and returned to baseline by 18 hours, whereas *BAZ1A* rose more gradually and remained elevated through 24 hours, consistent with a priming function for the lncRNA in the early damage response. In the clinical context, we propose that the mechanical and ischemic trauma of saphenous vein harvesting^33,34^ triggers DNA damage in medial VSMCs, activating the *BAZ1A-AS1/BAZ1A* axis and initiating the proliferative transcriptional program that underlies NP. This model is supported by the observation that DNA damage markers and cellular senescence accumulate in VSMCs during dedifferentiation and NP,^11,35^ and by the upregulation of *BAZ1A* in independent cohorts of pulmonary arterial hypertension and carotid atherosclerosis, suggesting a conserved role across vascular disease states.^23,24^

The ChIRP-qPCR experiments demonstrated an unexpected dissociation between the DNA- and RNA-level interactions of *BAZ1A-AS1*. At the chromatin level, *BAZ1A-AS1* bound selectively to the 3′ distal exons of *BAZ1A*, beyond the region of direct antisense complementarity, but interacted negligibly with neighboring loci including *SRP54* and *NFKBIA*, demonstrating highly specific chromatin engagement. In contrast, RNA pulldown demonstrated selective interaction with *CNN1*, *CCND1*, and *IL6* mRNAs, representing contractile, proliferative, and inflammatory transcriptional programs, but not with *BAZ1A* mRNA itself. This dissociation suggests that *BAZ1A-AS1* employs mechanistically distinct modes of action: a cis-regulatory function at the *BAZ1A* locus, and a separate capacity for selective mRNA engagement. Precedent for such dual functionality exists among natural antisense lncRNAs; *MEG3*, for example, forms RNA-DNA triplex structures within GA-rich sequences to regulate TGF-β pathway genes,^36^ and other lncRNAs have been shown to scaffold protein complexes at chromatin while simultaneously influencing mRNA stability or translation.^37,38^ The relatively low copy number of *BAZ1A-AS1* (∼20 per cell) suggests that these interactions are confined to specific genomic or subcellular loci rather than mediating broad transcriptomic reprogramming, consistent with the focused nature of the observed transcriptional changes.

Knockdown of *BAZ1A* in HSVSMCs reduced *BAZ1A-AS1* expression by approximately 50%, suggesting bidirectional regulation within the lncRNA-gene pair. Several mechanisms may account for this observation. The proximity of the *BAZ1A-AS1* promoter to the *BAZ1A* locus suggests that transcriptional disruption at *BAZ1A* could alter the local chromatin environment or transcriptional machinery availability for *BAZ1A-AS1*. Alternatively, shared regulatory elements may coordinate expression of both transcripts, such that silencing one impairs the other. Finally, a feedback mechanism may exist in which sustained *BAZ1A* transcription is required to maintain *BAZ1A-AS1* expression, consistent with the transient kinetics of *BAZ1A-AS1* induction relative to the more sustained *BAZ1A* response following UV exposure. Distinguishing among these possibilities will require promoter deletion and enhancer mapping studies.

*Baz1a* haploinsufficiency in mice recapitulated key features of the *in vitro* phenotype. *Baz1a*⁺/⁻ mice exhibited significantly attenuated neointima formation, reduced Ki67⁺ VSMC proliferation, and increased αSma expression following carotid artery ligation, accompanied by upregulation of *Acta2* and *Myh11* and downregulation of *Ccnd1* in ligated vessels. Notably, elastin breaks were increased, and elastin area fraction was reduced at 200 μm in *Baz1a*⁺/⁻ mice despite reduced neointima formation, while collagen content was unchanged, suggesting that the protective effect is consistent with reduced VSMC proliferative and migratory capacity observed *in vitro* rather than changes in collagen deposition. An intriguing observation was the paradoxical increase in *Ccnd1* and *Cdk4* expression in uninjured contralateral carotid arteries of *Baz1a*⁺/⁻ mice, despite no differences in contractile gene expression. Given the low baseline proliferative activity of these transcripts (CT ∼30-31), the low proliferative activity in quiescent vessels, and the use of a global knockout model, these changes may reflect contributions from non-VSMC cell types, and their functional significance remains unclear. Although *BAZ1A-AS1* is human-specific and lacks a murine ortholog, the vascular phenotype of *Baz1a*⁺/⁻ mice supports BAZ1A as a functional effector of the axis and demonstrates that even partial reduction is sufficient to confer vascular protection.

This study has several limitations. First, the *ex vivo* NP model lacks hemodynamic flow and circulating blood cells which represents a key biological limitation, as SV grafts are abruptly exposed to arterial pressure, pulsatility, and shear stress after implantation. Thus, further studies using validated ex vivo SV perfusion systems are required to confirm our data in the future. However, using this static culture model, we recently reported that *ex vivo* NP in HSV differs according to body mass index in men, with less NP in veins from overweight and moderately obese men compared to lean counterparts,^11^ a finding that parallels clinical observations in patients undergoing CABG.^39,40^ Second, global, rather than VSMC-specific *Baz1a* knockout mice were used, precluding definitive assignment of the vascular phenotype to VSMC-intrinsic mechanisms. Third, the carotid artery ligation model addresses arterial injury, and its relevance to venous graft pathophysiology is indirect. Fourth, *BAZ1A-AS1* is human-specific and lacks a murine ortholog, limiting *in vivo* assessment of the lncRNA component of the axis. Future studies using humanized knock-in mice or transgenic mice expressing human BAZ1A-AS1 are required to confirm the role of BAZ1A-AS1 in vivo. Fifth, although knockdown of *BAZ1A-AS1* in HSVSMCs and *BAZ1A* in both HSVSMCs and RD cells produced concordant phenotypic effects, direct comparison between experiments is limited by differences in knockdown methodology (LNA gapmeR ASO versus siRNA), cell types, and culture conditions. Finally, the precise molecular mechanisms by which *BAZ1A-AS1* engages its DNA and mRNA targets, and whether these interactions occur simultaneously, sequentially, or in distinct subcellular compartments, remain to be defined.

In conclusion, we identify *BAZ1A-AS1* as a stress-responsive regulator of VSMC dedifferentiation and NP, operating through a dual mechanism involving cis-regulatory chromatin interaction at the *BAZ1A* locus and selective engagement of phenotype-relevant mRNAs. These findings position the *BAZ1A-AS1/BAZ1A* axis as a potential therapeutic target for preventing saphenous vein graft failure following coronary artery bypass surgery and warrant further investigation into the precise molecular mechanisms governing this regulatory module.

## Supporting information

Supplemental Figures 1-5, Tables 1-4, uncroped blot images

## Acknowledgements

The authors acknowledge the Electron Microscopy and Histology Core Facility at Augusta University (RRID: SCR_026810) for tissue processing and histological services. This study was funded by grants NIH AG076235 (N.L.W), AHA 971459 (N.L.W), AHA 863622 (N.L.W), AHA 26BTPA1622831 (N.L.W, https://doi.org/10.58275/AHA.26BTPA1622831.pc.gr.243735) and AHA 23PRE1026496 (D.K).

