## Supplemental Figures 1-5, Tables 1-4, uncroped blot images for "A regulatory role of novel long non-coding RNA, *BAZ1A-AS1*, in vascular smooth muscle cell functions during neointima proliferation in human saphenous veins"

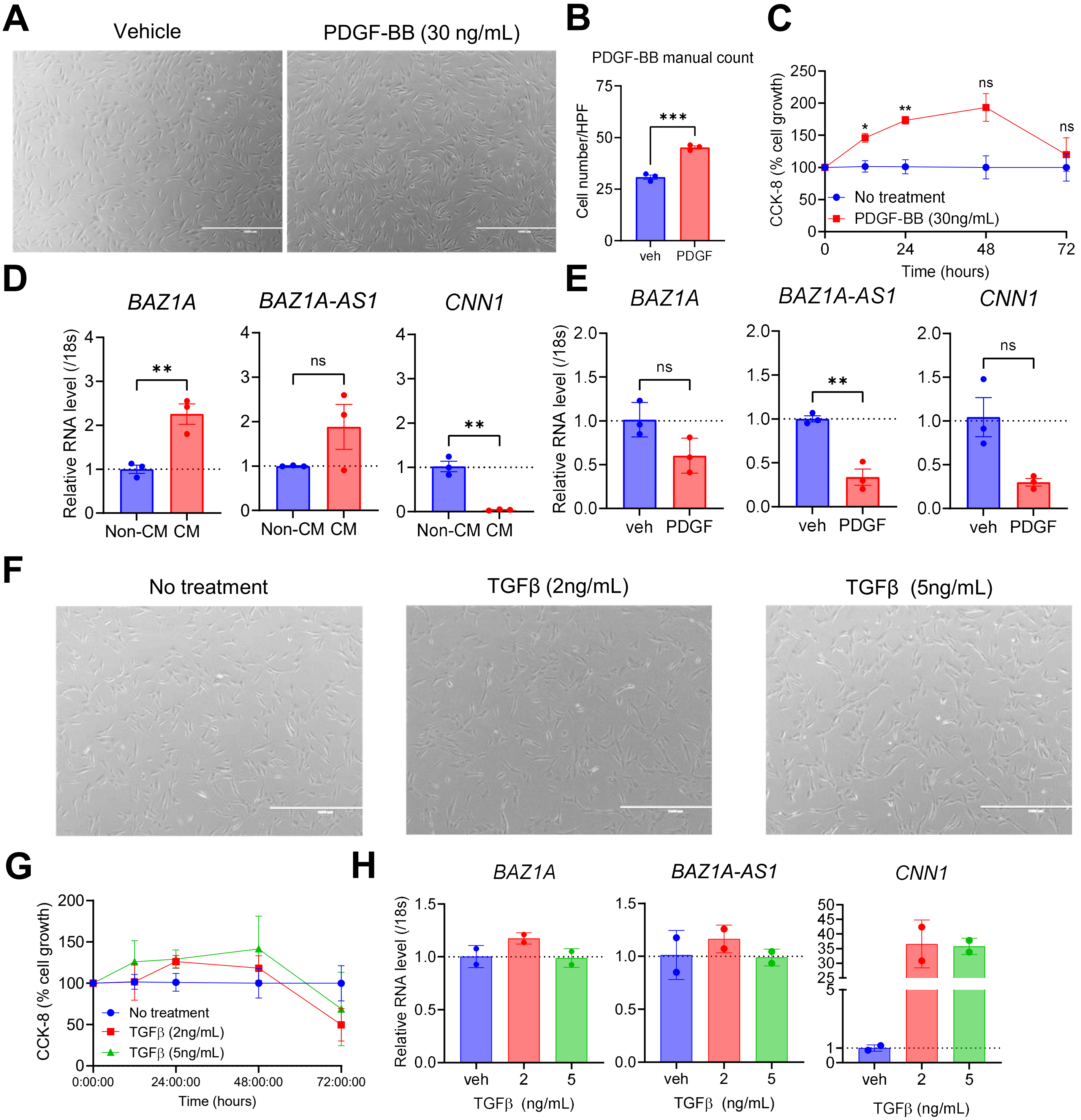


**Supplemental Figure S1.** *BAZ1A-AS1* and *BAZ1A* expression under growth factor and paracrine stimulation in HSVSMCs

**A**, Representative brightfield images of HSVSMCs treated with vehicle or PDGF-BB (30 ng/mL) for 48 h. Scale bar = 1000 μm. **B**, Manual cell count per high-power field (HPF) in vehicle versus PDGF-BB–treated HSVSMCs (n = 3). **C**, CCK-8 cell growth assay over 72 h in untreated and PDGF-BB–treated (30 ng/mL) HSVSMCs (n = 3). **D**, RT-qPCR of *BAZ1A*, *BAZ1A-AS1*, and *CNN1* in HSVSMCs treated with non-conditioned control media (Non-CM) or Day 7 HSV-conditioned media (CM) for 24 h (n = 3). **E**, RT-qPCR of *BAZ1A*, *BAZ1A-AS1*, and *CNN1* in HSVSMCs treated with vehicle (veh) or PDGF-BB (10 ng/mL) for 24 h (n = 3). **F**, Representative brightfield images of HSVSMCs treated with vehicle, TGFβ (2 ng/mL), or TGFβ (5 ng/mL) for 48 h. Scale bar = 1000 μm. **G**, CCK-8 cell growth assay over 72 h in HSVSMCs treated with vehicle, TGFβ (2 ng/mL), or TGFβ (5 ng/mL) (n = 3). **H**, RT-qPCR of *BAZ1A*, *BAZ1A-AS1*, and *CNN1* in HSVSMCs treated with vehicle (veh), TGFβ (2 ng/mL), or TGFβ (5 ng/mL) for 24 h (n = 3). Data presented as mean ± SEM. Statistical tests: two-tailed unpaired *t*-test (**B**, **D**, **E**); two-way ANOVA with Tukey post-hoc (**C**, **G**); one-way ANOVA with Dunnett's post-hoc vs vehicle (**H**). *$P < 0.05$, **$P < 0.01$, ***$P < 0.001$; ns, not significant.

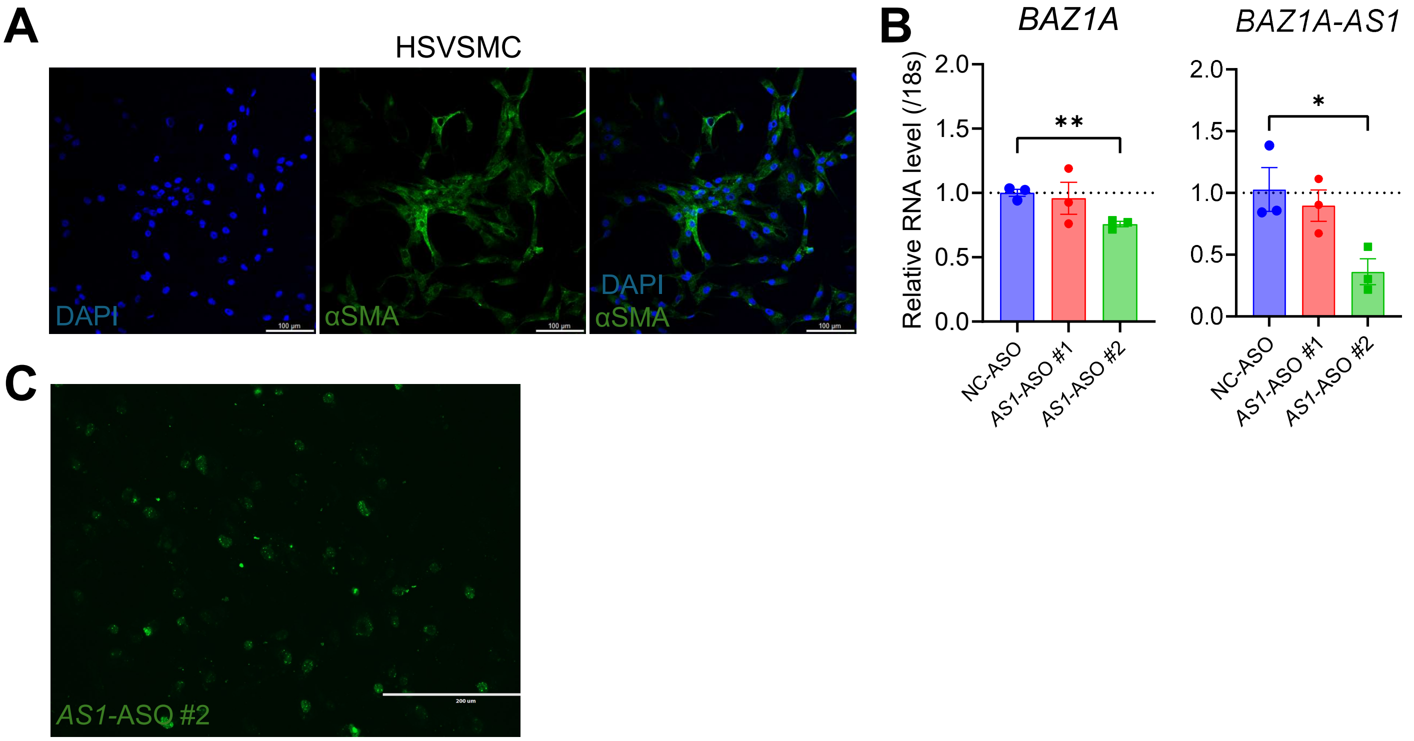


**Supplemental Figure S2.** Validation of LNA gapmeR antisense oligonucleotide knockdown efficiency in HSVSMCs. **A**, Representative immunofluorescence of HSVSMCs co-stained for DAPI (**blue**) and αSMA (**green**), confirming VSMC identity. Scale bar = 100 µm. **B**, RT-qPCR of *BAZ1A* (**left**) and *BAZ1A-AS1* (**right**) in HSVSMCs transfected with NC-ASO, *AS1*-ASO #1, or *AS1*-ASO #2 (n = 3). *AS1*-ASO #2 significantly reduced both *BAZ1A* and *BAZ1A-AS1* expression and was selected for all downstream experiments. **C**, Representative fluorescence image of FAM-labeled *AS1*-ASO #2 uptake in HSVSMCs. Scale bar = 200 µm. Data presented as mean ± SEM. Statistical test: one-way ANOVA with Dunnett's post-hoc vs NC-ASO (**B**). *$P < 0.05$, **$P < 0.01$, ***$P < 0.001$; ns, not significant.

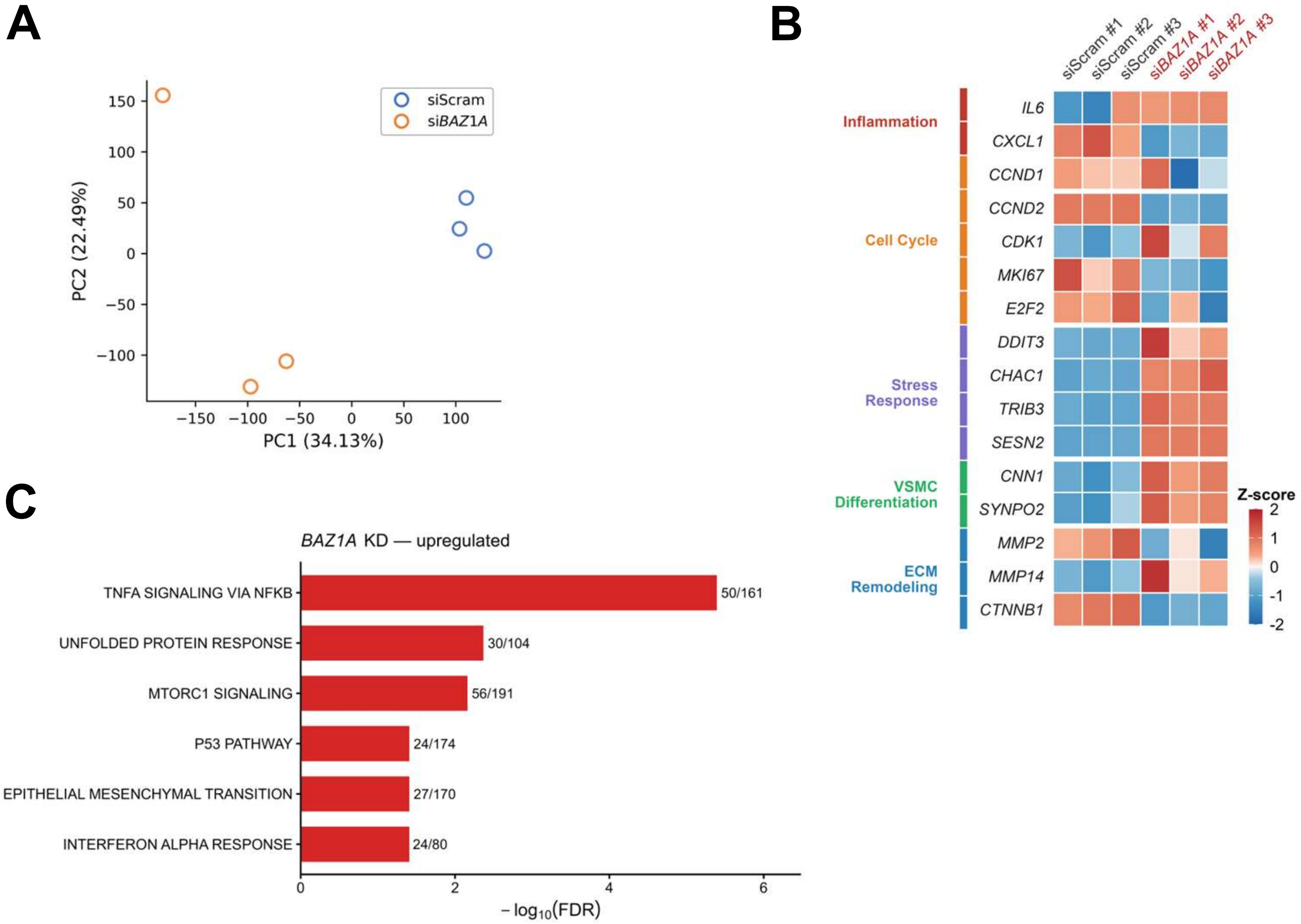


**Supplemental Figure S3.** Transcriptomic profiling of *BAZ1A* knockdown in RD cells under UV exposure
**A**, Principal component analysis of bulk RNA-seq in RD cells transfected with siScram or si*BAZ1A* under UV exposure (n = 3 per group). **B**, Heatmap of z-scored expression of representative genes across five functional categories (Inflammation, Cell Cycle, Stress Response, VSMC Differentiation, ECM Remodeling) in siScram versus si*BAZ1A*-treated RD cells (n = 3 per group). **C**, Hallmark gene set enrichment analysis (GSEA) of the *BAZ1A* knockdown transcriptome showing significantly upregulated pathways (FDR < 0.05), ranked by −log10(FDR). Bar labels indicate the number of leading-edge genes over total pathway gene set size. No Hallmark pathways reached significance (FDR < 0.05) in the downregulated direction, although E2F Targets and Reactive Oxygen Species Pathway showed non-significant trends (FDR = 0.096 each), consistent with proliferation suppression observed in the *BAZ1A-AS1* knockdown transcriptome (**Figure 5D**). The *BAZ1A* KD volcano plot is shown in main **Figure 5B** (right panel). Statistical tests: pre-ranked GSEA with 1,000 permutations and Benjamini-Hochberg FDR correction (C).

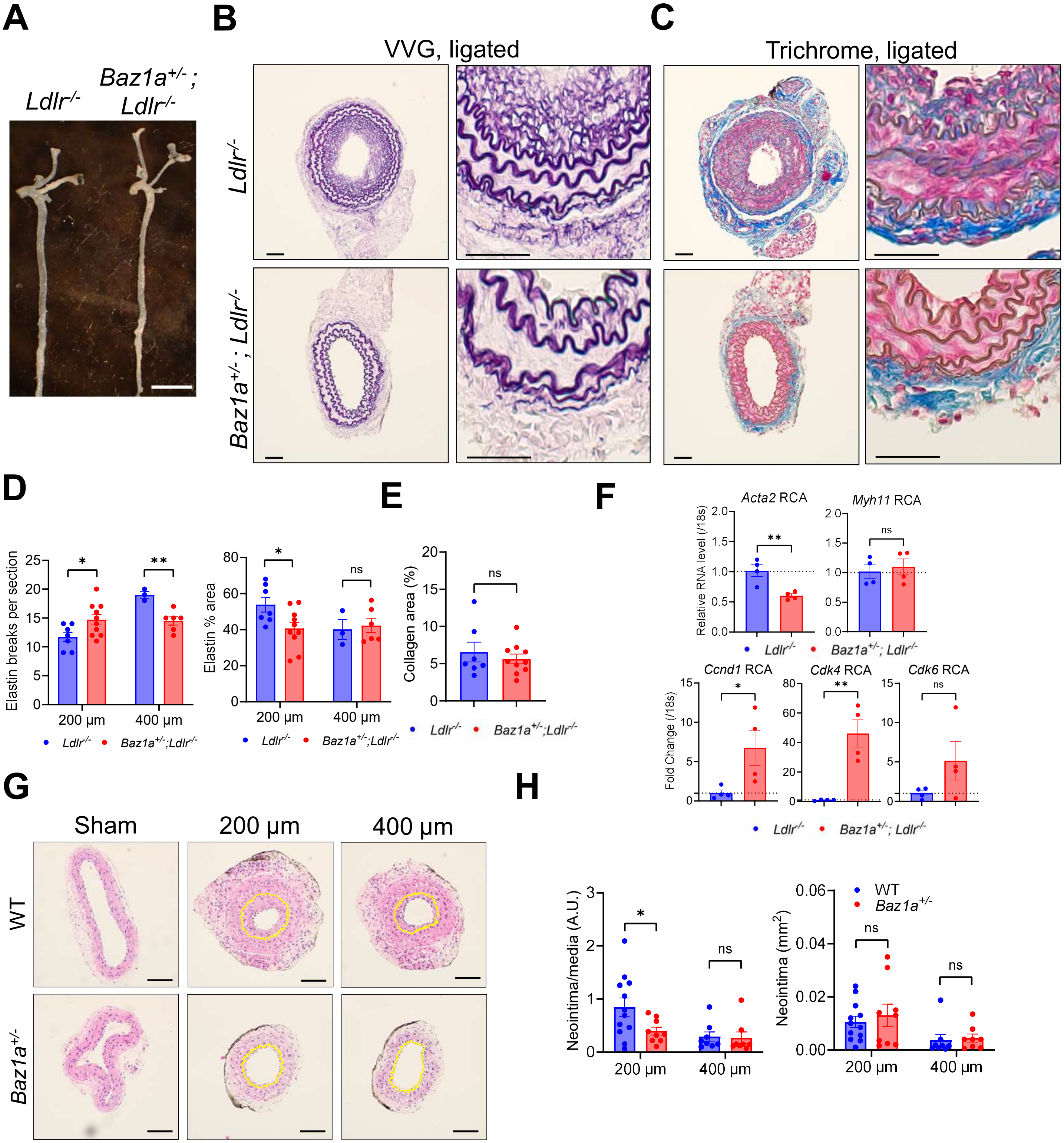


**Supplemental Figure S4.** Extended histological, morphometric, and gene expression analysis of carotid arteries after ligation surgery

**A**, Representative gross images of carotid bifurcations from *Ldlr*⁻/⁻ and *Baz1a*⁺/⁻;*Ldlr*⁻/⁻ mice 4 weeks after complete ligation. Scale bar: 5 mm. **B**, Representative Verhoeff-Van Gieson (VVG) staining of ligated carotid arteries from *Ldlr*⁻/⁻ and *Baz1a*⁺/⁻;*Ldlr*⁻/⁻ mice. Scale bars: 100 µm (cross-sections), 50 µm (insets). **C**, Representative Masson's trichrome staining of ligated carotid arteries from *Ldlr*⁻/⁻ and *Baz1a*⁺/⁻;*Ldlr*⁻/⁻ mice. Scale bars: 100 µm (cross-sections), 50 µm (insets). **D**, Quantification of elastin breaks per section (**left**) and elastin area fraction (**right**) in ligated carotid arteries at 200 µm and 400 µm proximal to the ligation site (n=3-10). **E**, Quantification of collagen area fraction by trichrome staining in ligated carotid arteries of *Ldlr*⁻/⁻ and *Baz1a*⁺/⁻;*Ldlr*⁻/⁻ mice (n=7-10). **F**, RT-qPCR of contractile genes *Acta2* and *Myh11* (**top**) and proliferation genes *Ccnd1*, *Cdk4*, and *Cdk6* (**bottom**) in uninjured right carotid arteries (RCAs) of *Ldlr*⁻/⁻ and *Baz1a*⁺/⁻;*Ldlr*⁻/⁻ mice (n = 4, 2 mice pooled per biological replicate). **G**, Representative H&E images of sham and ligated carotid arteries at 200 µm and 400 µm from WT and *Baz1a*⁺/⁻ mice (C57BL/6 background). Yellow dashed lines indicate internal elastic lamina. Scale bar: 100 µm. **H**, Neointima-to-media ratio (**left**) and neointima area (**right**) in WT and *Baz1a*⁺/⁻ mice at 200 µm and 400 µm proximal to the ligation site (n = 8–12). Data are presented as mean ± SEM. *$P < 0.05$, **$P < 0.01$; ns, not significant. Statistical comparisons by two-tailed unpaired t-test (**F**) or unpaired t-test/Mann-Whitney test as appropriate after Shapiro-Wilk normality testing (**D**, **E**, **H**).

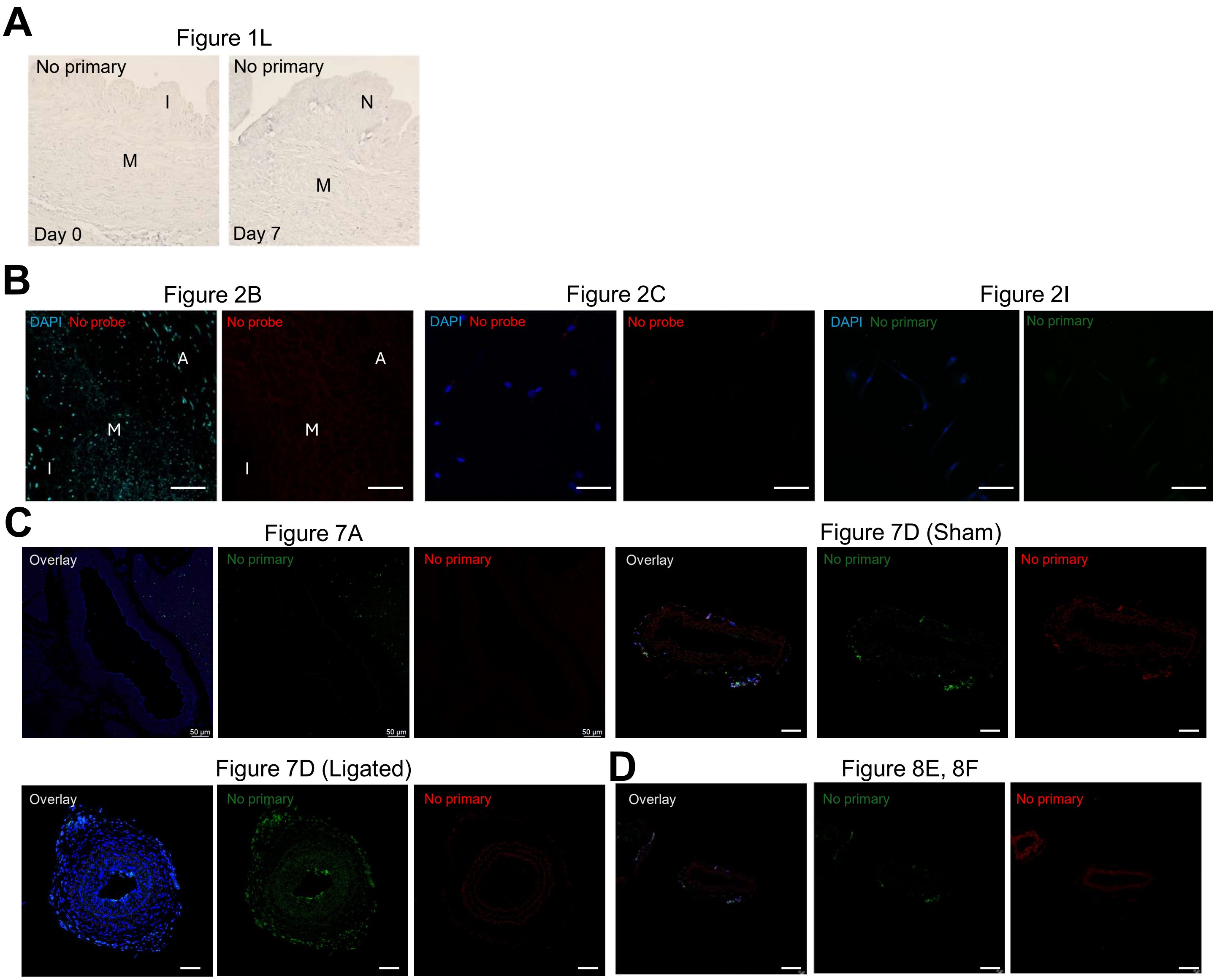


**Supplemental Figure S5**. Negative controls for immunohistochemistry, RNA-FISH, and immunofluorescence experiments

**A**, No-primary antibody controls for DAB immunohistochemistry of Day 0 (left) and Day 7 (right) HSV (corresponding to **Figure 1L**). I, intima; M, media; N, neointima. **B**, No-probe controls for Stellaris RNA-FISH of *BAZ1A-AS1* in Day 7 HSV (left two panels, corresponding to **Figure 2B**; scale bar: 100 µm) and in cultured HSVSMCs (middle two panels, corresponding to **Figure 2C**; scale bar: 50 µm). No-primary antibody control for BAZ1A immunofluorescence in HSVSMCs (right two panels, corresponding to **Figure 2I**; scale bar: 50 µm). **C**, Overlay images (left) and corresponding no-primary antibody controls (middle, right) for immunofluorescence staining of Baz1a and αSma in mouse aorta (left set, corresponding to **Figure 7A**) and carotid artery (right set, corresponding to **Figure 7D**). Scale bars: 50 µm. **D**, Overlay images (left) and corresponding no-primary antibody controls (middle, right) for immunofluorescence staining of Ki67 or Baz1a and αSma in ligated carotid arteries (corresponding to **Figure 8E-F**). Scale bars: 50 µm.

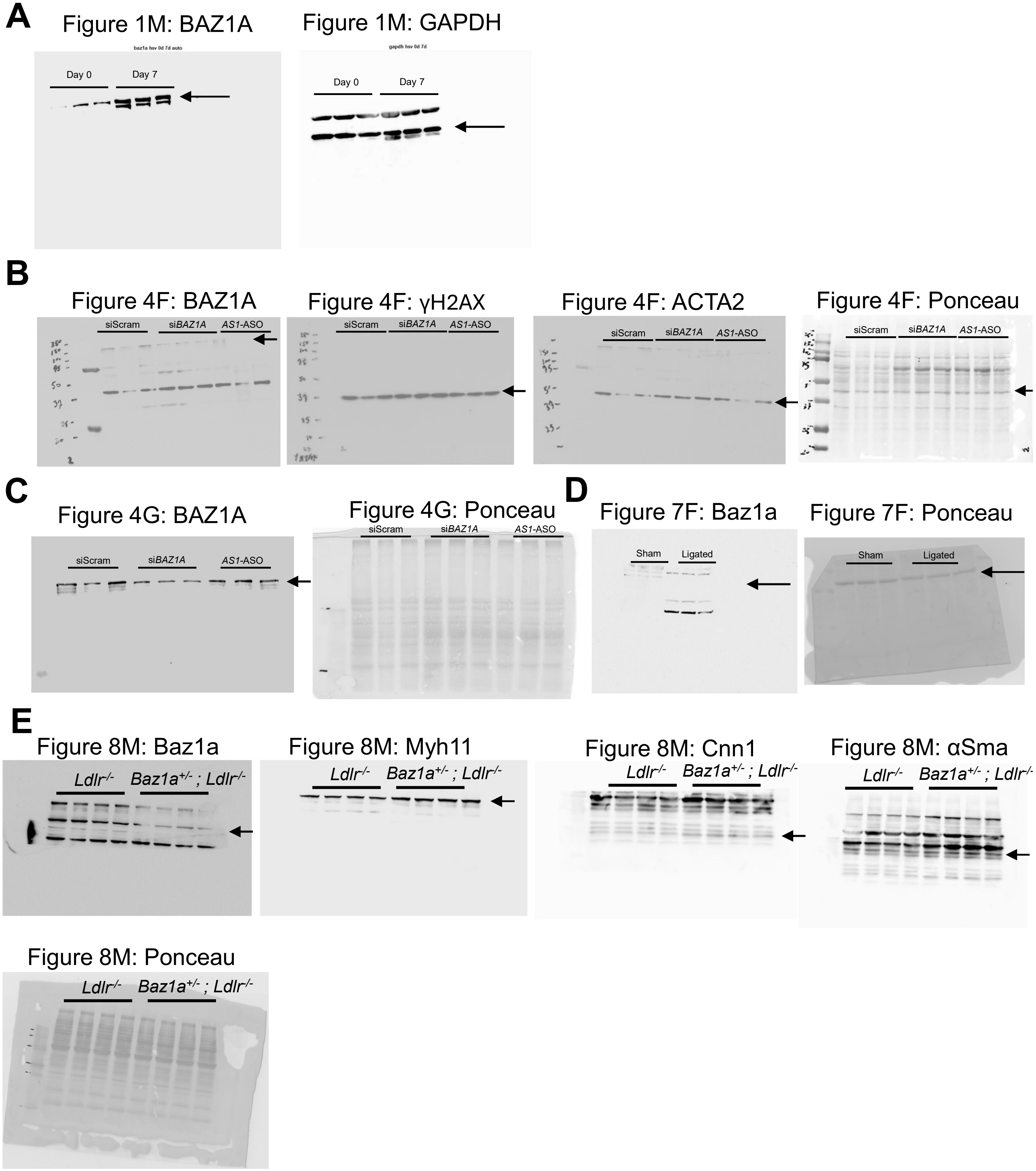


**Supplemental Figure S6.** Uncropped Western blot images

| Patient | Sex | Age | Weight (kg) | Height (cm) | BMI | EF (%) | DM | HF | HTN | DLD | Smoking | CKD | Other |
|---|---|---|---|---|---|---|---|---|---|---|---|---|---|
| 1 | M | — | 91.0 | 178 | 28.7 | 55 | No | No | Yes | Yes | Yes | No | — |
| 2 | M | — | 71.7 | 165 | 26.3 | 41 | No | No | Yes | Yes | Yes | No | — |
| 3 | M | — | 100.0 | 188 | 28.3 | 60–65 | Yes | No | Yes | Yes | No | No | — |
| 4 | F | — | 67.3 | 160 | 26.3 | 60 | Yes | No | Yes | Yes | Yes | No | — |
| 5 | M | — | 77.0 | 177 | 24.6 | 55–60 | No | No | Yes | Yes | No | No | Hx of malignancy |
| 6 | F | — | 86.0 | 165 | 31.6 | 60–65 | No | No | Yes | Yes | No | No | Hx of stroke/TIA |
| Summary | 4M/2F | — | 82.2 ± 12.4 | 172 ± 10 | 27.6 ± 2.4 | — | 2/6 (33%) | 0/6 (0%) | 6/6 (100%) | 6/6 (100%) | 3/6 (50%) | 0/6 (0%) | — |

**Supplemental Table S1.** Clinical characteristics of patients whose HSV were used for bulk RNA sequencing.
Values are presented as mean ± SD or n/N (%). All saphenous veins were harvested using the endoscopic vein harvesting (EVH) technique. BMI, body mass index; DM, diabetes mellitus; HF, heart failure; EF, ejection fraction; HTN, hypertension; DLD, dyslipidemia; CKD, chronic kidney disease; TIA, transient ischemic attack.

**Supplemental Table S2.** Primer sequences

| Species | Gene | Forward primer | Reverse primer |
|---|---|---|---|
| Human | BAZ1A-AS1 | AGGAGCTTTTCACTCCTCGC | TTATGAACCCCCAGCGCTTT |
| Human | BAZ1A #1 | GTGAAGAAACTGTCAAGCACCTC | GAACTCCTGCTTTGCCTGTCG |
| Human | MALAT1 215 | GAGCATATAATAATTCCAGGCACA | ATATACAATCAAGTCAAGCTCCTGAC |
| Human | MRGPRF | GGCTTCTCCATCAAGAGGAACC | TCAGGATGGAGAACACCGCCTT |
| Human | MRGPRF-AS1 | GGCCTACATGGCCCATACTG | CAGTGTCCAGTGCTTTGCAG |
| Human | GUCY1A1 | GCTCTTCTCAGACATCGTTGGG | ATAGGCATCGCCAATGGTCTCC |
| Human | GUCY1A1-AS1 | GCTGCCGTCTTCCCTGAATA | CCCAAGCTCTGGCAGTACAT |
| Human | CNN1 | ATGTCCTCTGCTCACTTCAAC | GCTGGTGGTCATACTTCTGG |
| Human | TAGLN | CATCCTGTCTGTCCGAACC | CACTATGATCCACTCCACCAG |
| Human | ACTA2 | TCAATGTCCCAGCCATGTAT | CAGCACGATGCCAGTTGT |
| Human | CCND1 | CATCTACACCGACAACTCCATC | TCTGGCATTTTGGAGAGGAAG |
| Human | CDK4 | GTGGCTGAAATTGGTGTCGG | GCCATCTGGTAGCTGTAGATTC |
| Human | CDK6 | GGCGCCTATGGGAAGGTGTTC | AAAGTCCAGACCTCGGAGAAGC |
| Human | NFKBIA | TGT GCT TCG AGT GAC TGA CC | TCA CCC CAC ATC ACT GAA CG |
| Human | SRP54 | GAA GTT GGG TTG TGG GGT CA | CTG AGT GCA CCA CCT CCT TT |
| Human | IL6 | ACTCACCTCTTCAGAACGAATTG | CCATCTTTGGAAGGTTCAGGTTG |
| Human | BAZ1A #2 | GGCGTGATCGCAGGGAAG | GGAGGTCTCCCTCTTCGTCT |
| Human | BAZ1A #3 | GCG CTG GGG GAA AAA TAC CT | AGA GGA ATT GGC CAA CTG GG |
| Human | BAZ1A #4 | GTC AGC AGG CCT GAC CAT AG | GGT TCC TGA ATA CCA CGC CA |
| Human | 18S | ATGGGCGGCGGAAAATAGC | TCTTGGTGAGGTCAATGTCTG |
| Human | GAPDH | CGACTTCAACAGCAACTCCCACTCTTCC | TGGGTGGTCCAGGGTTTCTTACTCCTT |
| Mouse | Baz1a | AAGCGAGAAGCGAAAAGCTC | AGCGTGAACGACGAGTAAGG |
| Mouse | Acta2 | CGAAACCACCTATAACAGCATCA | GCGTTCTGGAGGGGCAAT |
| Mouse | Cnn1 | ATGTCCTCTGCTCACTTCAAC | GCTGGTGGTCATACTTCTGG |
| Mouse | Myh11 | ACGACAACTCCTCACGATTC | TCACTTCTCATCTTCTCCTTGG |
| Mouse | Ccnd1 | GCGTACCCTGACACCAATCTC | CTCCTCTTCGCACTTCTGCTC |
| Mouse | Cdk4 | ATGGCTGCCACTCGATATGAA | TCCTCCATTAGGAACTCTCACAC |
| Mouse | Cdk6 | GGCGTACCCACAGAAACCATA | AGGTAAGGGCCATCTGAAAACT |
| Mouse | 18s | GCAATTATTCCCCATGAACG | GGCCTCACTAAACCATCCAA |

| Probe # | Probe Name | Sequence (5'→3') | Pool | 3' Modification |
|---|---|---|---|---|
| 1 | BAZ1A-AS1_1 | GACTTGAAAGTGACGGGTGG | Odd | Biotin |
| 2 | BAZ1A-AS1_2 | CCGACGAGGAAGTTTTCTAC | Even | Biotin |
| 3 | BAZ1A-AS1_3 | CCAATTTTGATAGGGAAGCG | Odd | Biotin |
| 4 | BAZ1A-AS1_4 | GGCGCGTCAGTCACACAAAA | Even | Biotin |
| 5 | BAZ1A-AS1_5 | CAGGACGAGAGGAGGTAGGG | Odd | Biotin |
| 6 | BAZ1A-AS1_6 | GGAACAAAGCGGAGCAGCTC | Even | Biotin |
| 7 | BAZ1A-AS1_7 | GATCAGAAACTTCCCAGGAG | Odd | Biotin |
| 8 | BAZ1A-AS1_8 | GTATCAGTGACAGGCTGTAA | Even | Biotin |
| 9 | BAZ1A-AS1_9 | CCTAGATCTTTTGGCTGATA | Odd | Biotin |
| 10 | BAZ1A-AS1_10 | CGTAATTGGAATCCCTTTTC | Even | Biotin |
| 11 | BAZ1A-AS1_11 | CTGATGTCAAGGTATCTTCT | Odd | Biotin |
| 12 | BAZ1A-AS1_12 | AGTTTCAAGAATGCTGGGGT | Even | Biotin |
| 13 | BAZ1A-AS1_13 | TGCTTTAGGGGAATTTTCGC | Odd | Biotin |
| 14 | BAZ1A-AS1_14 | CGTTCTGAGTTACTTCTAGT | Even | Biotin |
| 15 | BAZ1A-AS1_15 | TATCATTTCGCAGTATTGGT | Odd | Biotin |
| 16 | BAZ1A-AS1_16 | ATGAACAGTTAAAGCCAGCC | Even | Biotin |

**Supplemental Table S3.** Biotinylated antisense oligonucleotide probe sequences for ChIRP-qPCR targeting *BAZ1A-AS1*.
*All probes were synthesized by Biosearch Technologies with 3' biotin modification and purified by reverse-phase chromatography (RPC). Probes were designed to target BAZ1A-AS1, excluding the region overlapping with BAZ1A exon 1. Odd-numbered probes (1, 3, 5, 7, 9, 11, 13, 15) were assigned to the Odd pool; even-numbered probes (2, 4, 6, 8, 10, 12, 14, 16) were assigned to the Even pool. Scale: 50 nmol per probe.*

**Supplemental Table S4.** Antibodies used for Western blot and Immunohistochemistry

| Primary Antibodies for Western Blot/Immunofluoresence | | | |
|---|---|---|---|
| **Antibody** | Vendor or Source | Catalog # | Working concentration |
| **BAZ1A/Baz1a** | Bethyl | A301-318A | 1:1000 |
| **BAZ1A** | Novus | NB100-61041 | 1:1000 |
| **yH2AX** | Sigma-Aldrich | 05-636-25UG | 1:1000 |
| **ACTA2** | Proteintech | 15117-1-AP | 1:500 |
| **αSMA/αSma** | Abcam | ab5694 | 1:1000 |
| **Ki67** | Abcam | ab15580 | 1:1000 |
| **NOS3** | BD | 612392 | 1:1000 |
